# Genetic screens in isogenic mammalian cell lines without single cell cloning

**DOI:** 10.1101/677385

**Authors:** Peter C DeWeirdt, Kendall R Sanson, Ruth E Hanna, Mudra Hegde, Annabel K Sangree, Christine Strand, Nicole S Persky, John G Doench

**Affiliations:** Genetic Perturbation Platform, Broad Institute of MIT and Harvard, Cambridge, MA

**Author notes:** These authors contributed equally (Peter C DeWeirdt, Kendall R Sanson, Ruth E Hanna).

## Abstract

Isogenic pairs of cell lines, which differ by a single genetic modification, are powerful tools for understanding gene function. Generating such pairs for mammalian cells, however, is labor-intensive, time-consuming, and impossible in some cell types. Here we present an approach to create isogenic pairs of cells and screen them with genome-wide CRISPR-Cas9 libraries to generate genetic interaction maps. We queried the anti-apoptotic genes BCL2L1 and MCL1, and the DNA damage repair gene PARP1, via 25 genome-wide screens across 4 cell lines. For all three genes, we identify a rich set of both expected and novel buffering and synthetic lethal interactions. Further, we compare the interactions observed in genetic space to those found when targeting these genes with small molecules and identify hits that may inform the clinical uses for these inhibitors. We anticipate that this methodology will be broadly useful to comprehensively study genes of interest across many cell types.

---

Genetic interaction networks can suggest functional roles of uncharacterized genes and capture subtle biological interactions, which may prove critical for interpreting genetic signal from genome-wide association studies of common disease states. Crosses of yeast knockout strains have yielded rich networks of genetic interactions, and have further shown that the shape of the network will change based on growth conditions^1–5^. In mammalian cells, the construction of such networks is orders of magnitude more complicated, due to increased genome size, the diversity of cell types, and numerous technical factors. One approach is to use either RNAi^6^ or CRISPR technology^7–11^ to screen a library of all possible combinatorial perturbations within a focused gene list. This approach has been used to generate genetic interaction maps for up to hundreds of genes^12^; however, screening all combinations of protein coding genes in the human genome would require, at bare minimum, approximately 400 million perturbations and 200 billion cells, which is equivalent to 5,000 concurrent genome-wide screens with typical guide libraries and currently exceeds the practical limits of tissue culture. This scale is exacerbated by the diversity of cell types in which to study such interactions.

A second, complementary approach to query genetic interactions leverages isogenic pairs of human cells, akin to mutant strains of model organisms, to enable the delineation of a given gene’s contribution to phenotypes of interest. Initial gene targeting approaches in human cell lines to create even a single knockout have yielded valuable insights but were quite laborious to generate^13–19^. Today, CRISPR technology has made cell line engineering possible for a broad range of researchers, but that is distinct from making it easy. Creating the site-specific nuclease for Cas9 is as simple as ordering a short nucleic acid, in contrast to the more expensive and time-consuming task of assembling a customized pair of zinc finger nucleases or TALENs^20^. After design of the targeted nuclease, however, substantial work remains: the isolation of single cells, often across multiple 96-well plates; culture of those cells for several weeks while colonies form; isolation of genomic DNA from replicated plates; and finally, PCR, sequencing, and analysis to determine which colonies have the intended genotype^21^. Indeed, off-the-shelf knockout clones, which are available in only a very limited number of cell lines, can be purchased from vendors for thousands of dollars, and the customized generation of a knockout clone in a cell line of interest costs tens of thousands of dollars. Thus, there is a great need for approaches that obviate the need to generate single cell clones and enable the creation of large scale genetic interaction maps for genes of interest in their appropriate cellular context.

Here, we leverage orthogonal Cas enzymes from *S. pyogenes* and *S. aureus* to conduct genome-wide CRISPR screens in paired mutant cell lines without the need for single cell cloning; we call this approach “anchor screening,” as the single genetic mutant “anchors” the resulting interaction network. We selected BCL2L1, MCL1, and PARP1 as anchor genes, as they each have well-established genetic interactions to allow benchmarking, and they are also the subject of intense clinical development, allowing both a comparison between small molecule inhibition and genetic knockout and, for PARP inhibitors, potentially an expansion of the genotypes beyond BRCA1 and BRCA2 mutant tumors in which these drugs may show efficacy. The rich set of uncovered genetic interactions shown here coupled with the ease of conducting such screens illustrate the power of this technology.

## RESULTS

Genetic screens with CRISPR technology often start with the creation of a cell line stably expressing Cas9, usually integrated into the genome via lentivirus or piggybac transposase^22, 23^. Because only a single element is delivered, this can be performed at small-scale, and the resulting cells expanded over the course of several weeks to the tens of millions of cells required for genome-scale libraries of single guide RNAs (sgRNAs, hereafter referred to as “guides”). In theory, one could also introduce a guide targeting a gene of interest at this step, to create a pool of knockout cells, and subsequently screen that population of cells against a library of guides. However, if there is any selective pressure against the knockout cells, they will be selected against during scale-up (Supplementary Fig. 1). For example, assume that i) unmodified cells, or those with in-frame indels, double every 24 hours, and ii) knockout cells represent 90% of the pool at the start; if the knockout cells have a 20% slower growth rate, they will represent less than half of the population after 3 weeks of proliferation. Inducible CRISPR systems could be helpful, but all of them require the use of additional components, such as recombinases, degrons, dimerization domains, transcriptional activators, or transcriptional repressors, as well as small molecule inducers, many of which have biological effects. Further, recent comparisons have shown that current systems often have substantially less activity than constitutive versions, or demonstrate leakiness; additionally, performance is typically cell-type dependent^24, 25^. Thus, there is a need for a simple method to generate cells poised for gene editing, expand them with no selective pressure, and trigger efficient knockout only when ready to begin a genetic screen.

**Figure 1.**
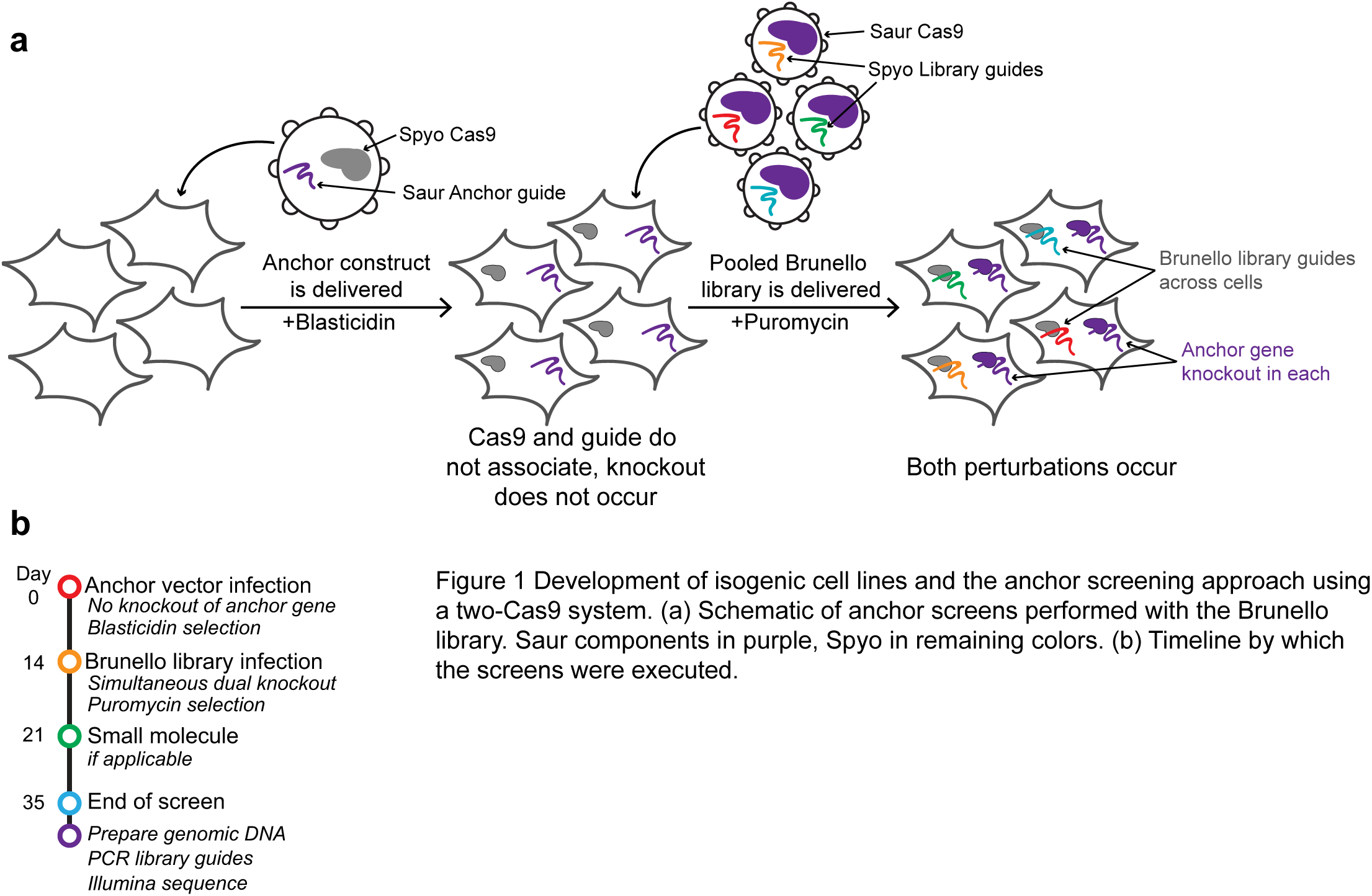
Development of isogenic cell lines and the anchor screening approach using a two-Cas9 system. (a) Schematic of anchor screens performed with the Brunello library. Saur components in purple, Spyo in remaining colors. (b) Timeline by which the screens were executed.

Previously, we and others developed *S. aureus* Cas9 (SaurCas9) for screening applications and paired it with *S. pyogenes* Cas9 (SpyoCas9) to enable combinatorial screens of “some-by-some” genes^10, 26^. We realized that small modifications to the vector designs would enable us to perform “one-by-all” screens with a workflow identical to standard genome-wide screens. The first vector, deemed the “anchor” vector, delivers SpyoCas9 and a guide compatible with *S. aureus* Cas9 (Saur-guide); the second “library” vector delivers SaurCas9 and a guide cassette compatible with *S. pyogenes* Cas9 (Spyo-guide), which is used to deliver the library of choice (Fig. 1a). With this approach, a guide targeting a gene of interest can be cloned into the anchor vector, delivered at small-scale, and the resulting population of cells expanded. Critically, because the guide is paired with the wrong Cas9, no editing will occur and thus there is no selective pressure during the cell scale-up. Finally, the library is introduced, and each cell will generate approximately simultaneous knockout of both the anchor gene and the gene targeted by the library without the need to validate any inducible components (Fig. 1b). This process can be completed in ∼5 weeks, which is less time than is required to generate and validate single cell clones, let alone screen them.

### Anchor screens for the anti-apoptotic genes BCL2L1 and MCL1

We selected two genes as anchors on which to test this approach, the anti-apoptotic genes MCL1 and BCL2L1, which themselves are a well-validated synthetic lethal pair^9, 10, 27^. For each gene we selected a previously-validated Saur-guide^10^ for use in the anchor vector and generated stable populations in the Meljuso melanoma cell line and the OVCAR8 ovarian cancer cell line; we also used the empty anchor vector to generate a control population. Into the library vector we introduced the Brunello genome-wide library, which has 4 guides per gene and ∼78,000 total guides^28^. We infected the library vector into the resulting 6 cell lines in duplicate, selected infected cells with puromycin for 5-7 days, and subsequently maintained the population with at least 500x coverage for an additional 2 weeks. As an additional experimental arm, we treated the control cells with either A-1331852 or S63845, small molecule inhibitors of BCL2L1^29^ or MCL1^30^, respectively, for the final 2 weeks of the experiment (Fig. 1b). At the end of the screen, we pelleted cells, prepared genomic DNA, retrieved the library guides by PCR, and performed Illumina sequencing to determine the abundance of each guide in each condition.

To detect genetic interactions with the anchor gene, we first calculated the log2-fold-change (LFC) compared to the initial library abundance, as determined by sequencing the plasmid DNA (Supplementary Data 1) and observed that replicates were well correlated (**Table 1**). For each anchor cell line, we then compared LFC values relative to the corresponding control cell line by fitting a nonlinear function and calculating the residual for each guide; a positive residual represents a buffering interaction, and a negative residual represents a synthetic lethal interaction (Fig. 2a). Residuals for individual guides were then averaged to determine a gene-level score, and statistical significance was determined by using a two-tailed Z-test (Fig. 2b); the same approach was used to determine sensitivity and resistance genes for the small molecules. The gene-level results from all screens are available in **Supplementary Data 2**.

**Figure 2.**
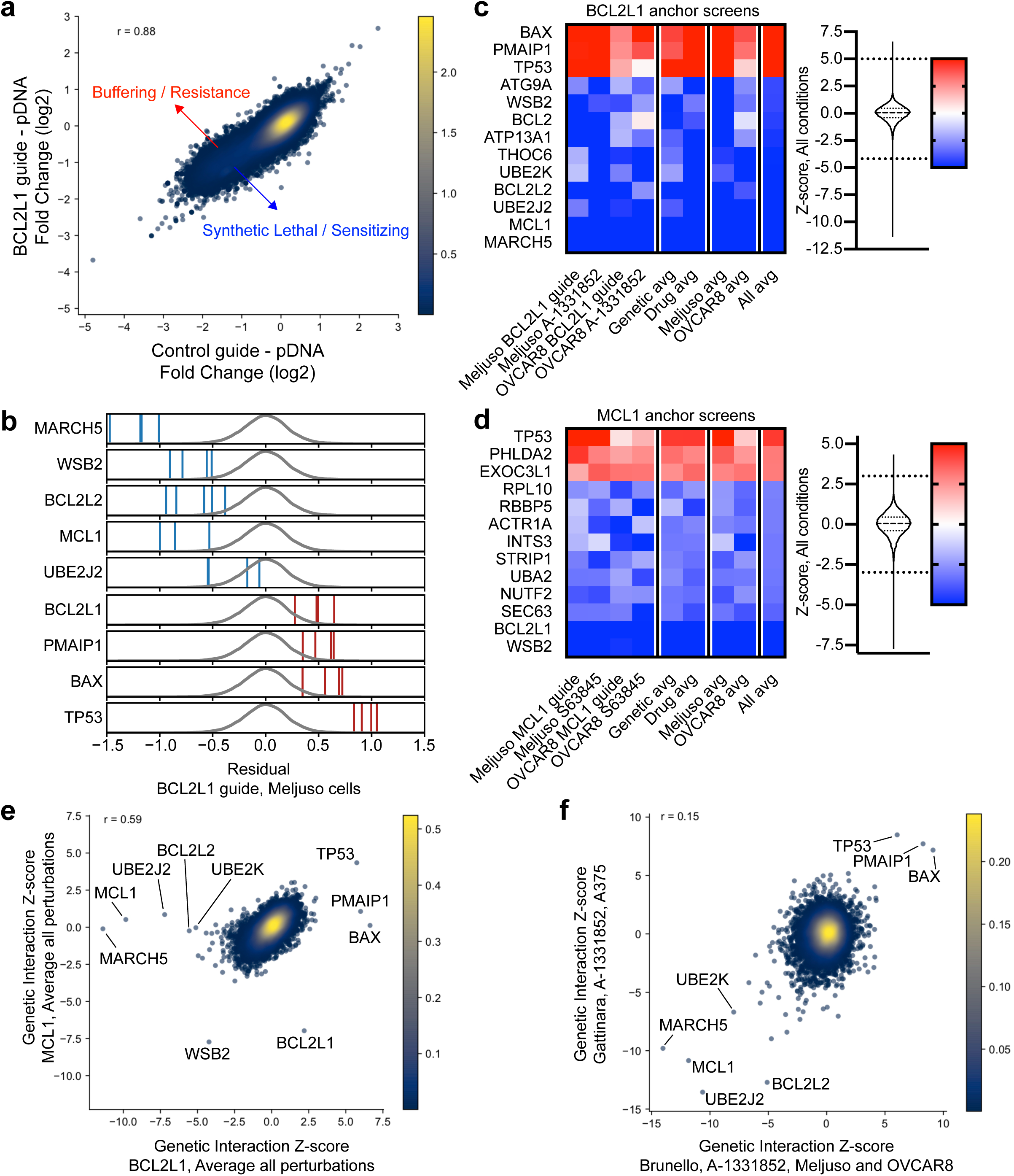
Anchor screens of BCL2L1 and MCL1 recover known and novel interactions. (a) Average log-fold changes for guides in Meljuso for control and BCL2L1 knockout lines. Points are colored by density. Pearson correlation coefficient is indicated. (b) Residuals for guides from the BCL2L1 anchor screen in Meljuso. Blue and red lines correspond to lethal and buffering guides respectively. Density of all guides is indicated by the grey distribution. (c) Top hits by average Z-score across all BCL2L1 screens. Color scale of Z-scores is shown to the right. A violin plot representing the distribution of Z-scores is adjacent to the color scale, along with two dotted lines representing the cutoffs for gene hits shown. The color scale is floored at −5 and ceilinged at 5. (d) Top hits by average Z-score across all MCL1 screens, as in (c). (e) Comparison of average Z-scores for all MCL1 and BCL2L1 perturbations screened with the Brunello library. (f) Z-scores for A-1331852 screened with Brunello and averaged for Meljuso and OVCAR8 cells vs Z-scores for A-1331852 screened with Gattinara in A375.

When anchoring on BCL2L1 knockout (Fig. 2c), MCL1 scored strongly in both Meljuso (rank 2, Z-score −9.6) and OVCAR8 (rank 2, Z-score −6.1). Conversely, when anchoring on MCL1 (Fig. 2d), BCL2L1 emerged as a top synthetic lethal interaction in both Meljuso (rank 5, Z-score −5.0) and OVCAR8 (rank 1, Z-score −7.3). These relationships were also captured by the parallel small molecule screens. With the BCL2L1 inhibitor A-1331852, MCL1 was a top sensitizer gene in both Meljuso (rank 2, Z-score −15.9) and OVCAR8 (rank 2, Z-score −7.8). Likewise, when screened with the MCL1 inhibitor S63845, BCL2L1 scored strongly in both Meljuso (rank 2, Z-score −9.7) and OVCAR8 (rank 1, Z-score −5.9). Thus, these genome-wide anchor screens were able to identify the expected synthetic lethal relationship between these genes, which were also observed with small molecule inhibition.

Other genes with well-established roles in apoptosis scored in these screens. Previously, we reported that BCL2L1 and BCL2L2 have a synthetic lethal relationship^10^, and that was borne out in these genome-wide screens: in the BCL2L1 anchor screen, BCL2L2 scored strongly in Meljuso (rank 4, Z-score −6.8) and OVCAR8 (rank 3, Z-score −5.2). Additionally, BCL2 scored highly in Meljuso (rank 3, Z-score −7.9) but was weak in OVCAR8 (rank 926, Z-score −1.9), a cell-type difference that we also observed previously^10^. Further, we saw strong buffering interactions between BCL2L1 and the pro-apoptotic genes TP53 (average rank 1, Z-score 5.4), BAX (rank 2, Z-score 4.2), and PMAIP1 (also known as NOXA, rank 3, Z-score 3.7). These genes were also the top 3 resistance hits for the small molecule A-1331852.

Additional genes emerged as strong hits in these screens (Fig. 2e). The E3 ubiquitin ligase MARCH5 showed strong synthetic lethality with BCL2L1 in both Meljuso (rank 1, Z-score −11.3) and OVCAR8 (rank 1, Z-score −6.1). Previous studies have shown that MCL1 levels are elevated in MARCH5 knockout cells^31^, and siRNA-mediated knockdown of MARCH5 led to loss of MCL1-mediated resistance to the BCL2-family inhibitor ABT-737^32^. Two additional top synthetic lethal hits with BCL2L1 are the E2 ligases UBE2J2 (rank 3, Z-score −7.2 across all conditions) and UBE2K (rank 5, Z-score −5.1). Another top-scoring gene was WSB2, a relatively unstudied gene that contains a SOCS box, a domain proposed to recruit ubiquitination factors to bound proteins^33^. This gene scored as a top synthetic lethal hit in the MCL1 anchor screens in both Meljuso (rank 2, Z-score −5.8) and OVCAR8 (rank 16, Z-score −4.7), as well as with the MCL1 small molecule inhibitor S63845 (rank 1, Z-score −14.5 in Meljuso; rank 2, Z-score −5.8 in OVCAR8). Thus, these screens connected several novel and understudied genes to the intrinsic apoptosis pathway via genetic evidence, and these are worthy of future biochemical study to determine their mechanism.

### Network analyses

To understand the generalizability of these novel relationships, we queried the Cancer Dependency Map (DepMap)^34, 35^, a compendium of genome-wide RNAi and CRISPR screens performed across hundreds of cancer cell lines. Here, correlation in fitness effects across cell lines suggests a functional relationship between genes^36–38^. Focusing on the CRISPR data screened with the Avana library, MARCH5 and MCL1 show a strong co-dependency (R = 0.66); UBE2J2 (R = 0.38) is the second-best correlate of MARCH5 dependency after MCL1, and UBE2K ranks 5th (R = 0.29). Furthermore, the best correlate to WSB2 essentiality is BCL2L2 (R = 0.39) and MCL1 is ranked 2nd (R = 0.29). Likewise, in the Project DRIVE RNAi screens^39^, the top correlate of WSB2 co-essentiality is MCL1 (R = 0.47), BCL2 ranks 4th (R = 0.39) and MARCH5 ranks 7th (R = 0.39).

Encouraged by these observations, we used a network approach to organize these data further. We selected the top 210 genes (nodes) with an absolute average Z-score greater than 2 across all BCL2L1 screens, and used co-essentiality correlations from the DepMap as edges, with an absolute cutoff of 0.2, which represent 0.45% of all correlations in the dataset (Supplementary Fig. 2). We used a graph based community detection algorithm^40^ to uncover clusters within the co-essentiality network (Supplementary Fig. 3). The clustering revealed densely connected groups of genes, one of which contained 9 of our top 12 hits by absolute average Z-score. In this cluster MCL1 and MARCH5 are connected with 10 and 9 genes respectively, making them the two most central hits of the group (Fig. 3a).

**Figure 3.**
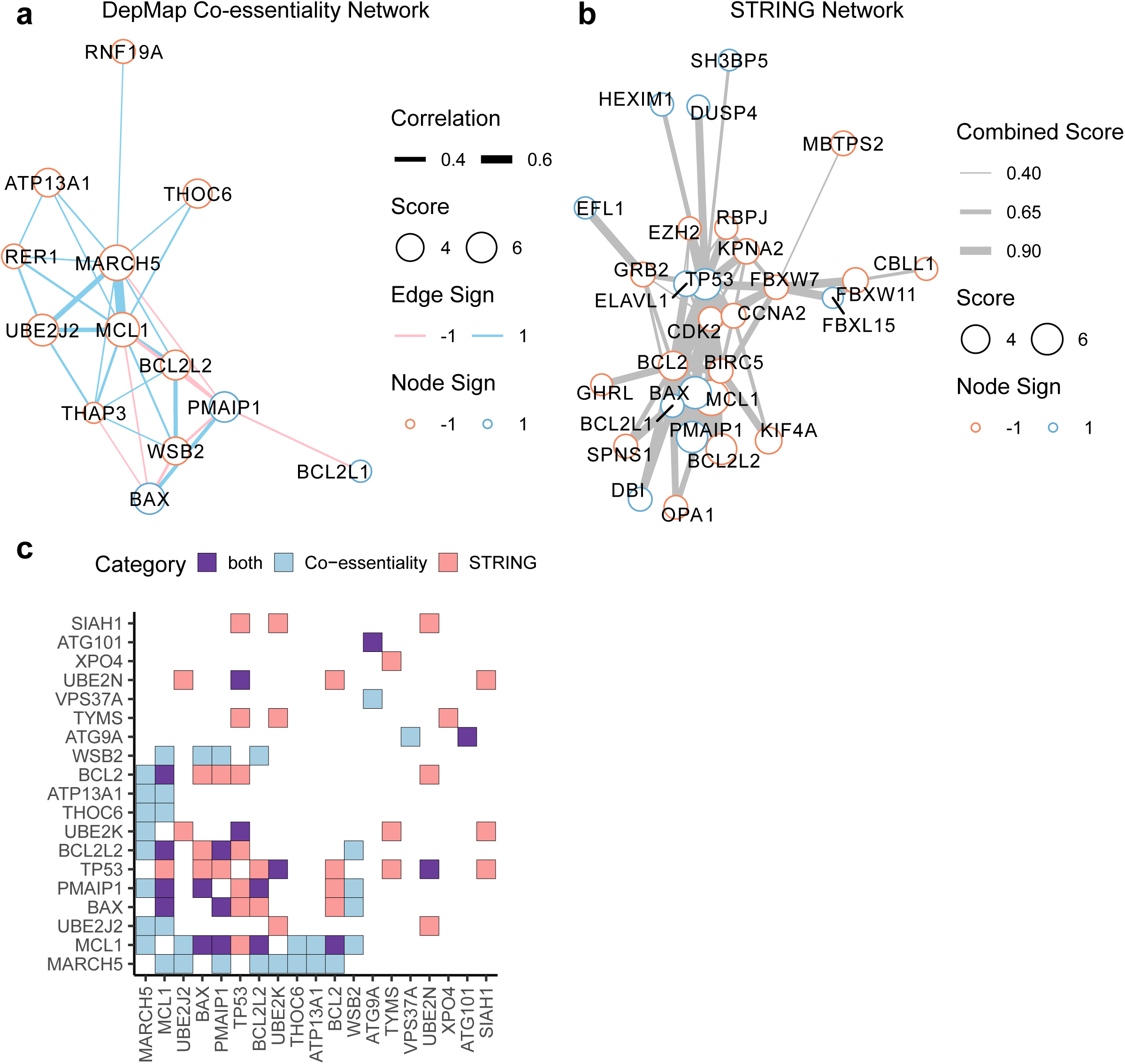
BCL2L1 anchor screens reveal functionally coherent clusters of genes. (a) Cluster of top hits from the DepMap co-essentiality network. Nodes represent genes and the size of each node is proportional to its average Z-score across all screens. Edges represent correlations in DepMap. (b) Cluster of top hits from the STRING network. Nodes are the same as (a). Edges represent combined score in STRING. (c) Interactions between the top 20 hits from both network sources. Genes are ordered by absolute average Z-score.

We also examined the STRING database^41^, which aims to build a global network of gene interactions based on protein-protein interactions, gene ontologies, and other curated annotations. We used a combined score cutoff of 400 to define edges in the STRING network (Supplementary Fig. 4), which corresponds to a medium confidence cutoff. We highlight one cluster that contained many of the strongest hits (Fig. 3b). In both the STRING and DepMap networks we saw an enrichment for edges when compared with random networks of genes of the same order (Supplementary Fig. 5a). Thus orthogonal network sources reveal a high level of connectivity between the top genes identified by these anchor screens.

**Figure 4.**
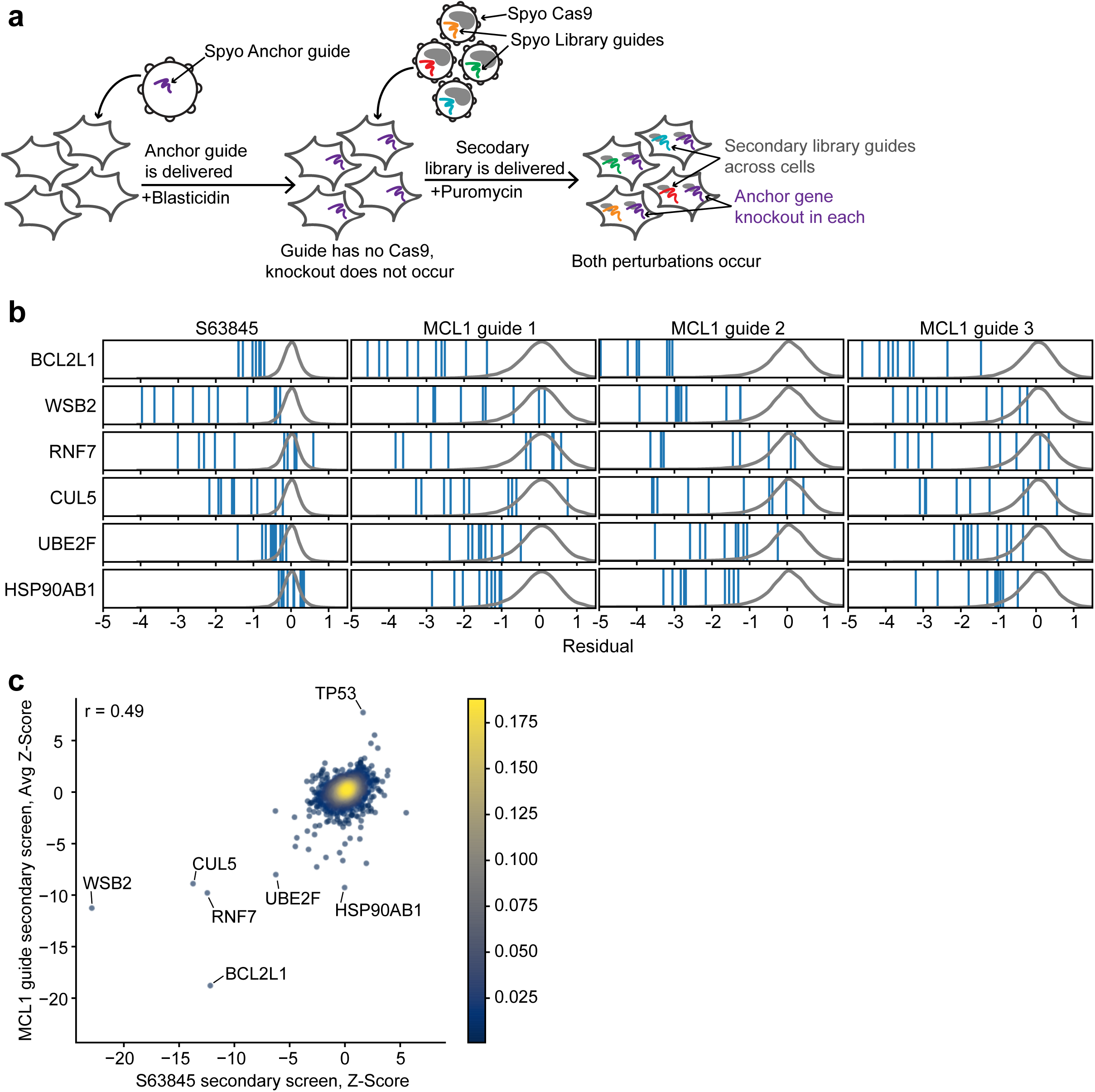
Alternate approach to anchor screens using a single Spyo-Cas9 system. (a) Schematic of Spyo-only approach screened with the secondary library utilizing only Spyo-Cas9. Spyo anchor perturbation shown in purple, Spyo library perturbations shown in remaining colors. (b) Residuals for guides from the secondary anchor screens in A375. Densities of all guides are indicated by the grey distributions. (c) Comparison of Z-scores for S63845 vs MCL1-knockout, averaged across all guides, screened with the secondary library in A375 cells. Points are colored by density. Pearson correlation coefficient is indicated.

**Figure 5.**
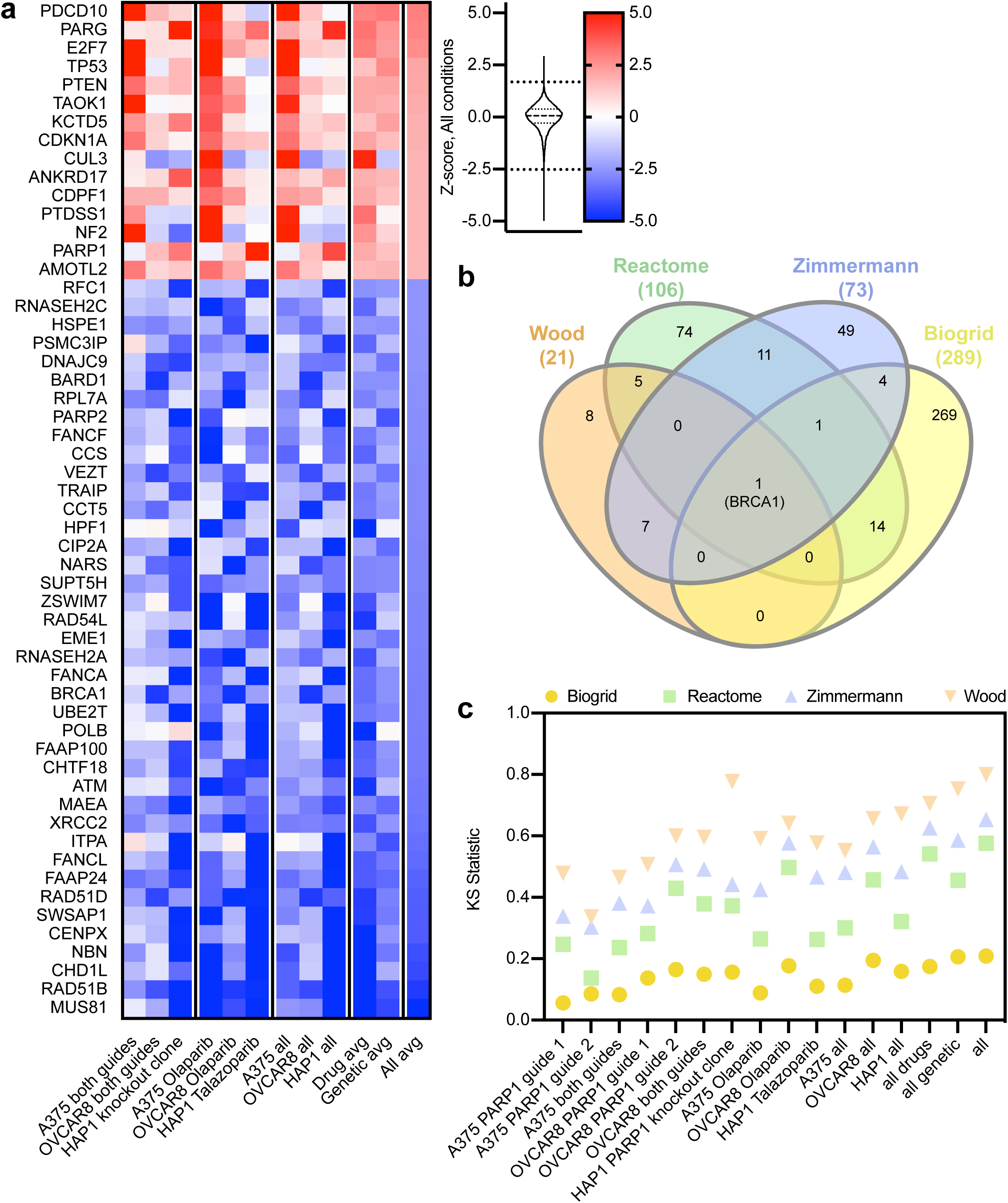
Top scoring genes from PARP screens are enriched in related gene sets. (a) Top hits by average Z-score across all PARP screens. Color scale of Z-scores is shown to the right. A violin plot representing the distribution of Z-scores is adjacent to the color scale, along with two dotted lines representing the cutoffs for gene hits shown. The color scale is floored at −5 and ceilinged at 5. (b) Venn diagram of curated gene sets included in the analysis. (c) KS statistic for each gene set. Statistic is shown for each screen and averages of various conditions. The value shown represents the alternative hypothesis that the cumulative distribution of genes in the gene set is greater than the distribution of genes not in the set.

Of the top 20 hits in the screen, all of them are connected to at least one other gene in at least one of the two networks, including 5 that are only detected in the DepMap network, showing that co-essentiality can reveal functional relationships that are currently unannotated in the STRING database (Fig. 3c). Finally, we performed the same analyses for the hits from the MCL1 anchor screen (Supplementary Fig. 5b, 6, 7). We again saw an enrichment for edges compared to random networks with both STRING and DepMap.

**Figure 6.**
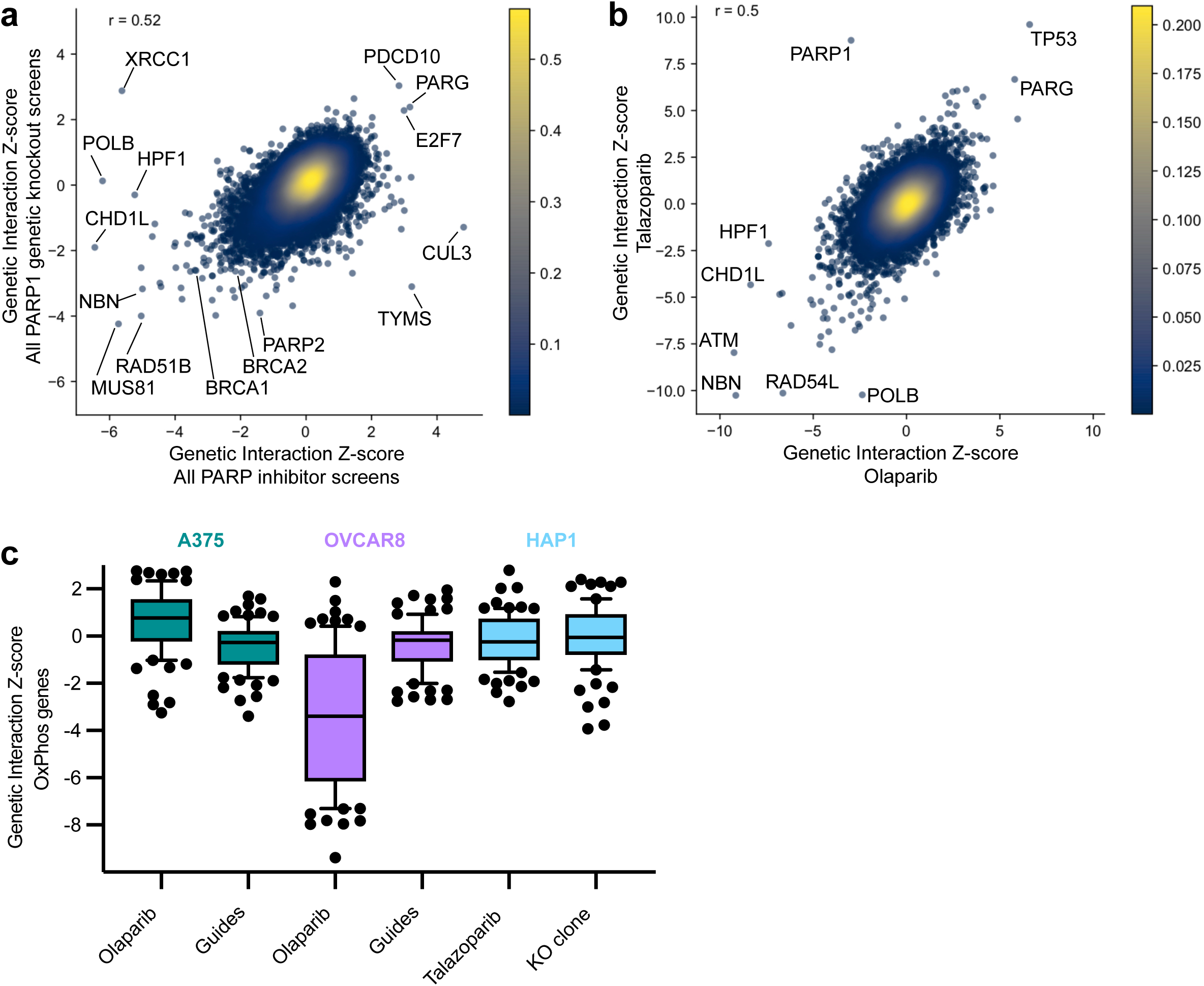
Hits from PARP screens agree across contexts with some notable exceptions. (a) Average Z-scores for PARP inhibitors and genetic knockout perturbations, screened with the Brunello library. Points are colored by density. Pearson correlation coefficient is included in the top left. (b) Z-scores for Olaparib and Talazoparib perturbations, screened with the Gattinara library. Points are colored by density. Pearson correlation coefficient is indicated. (c) Box plots of Z-scores for genes in the GO gene set “oxidative phosphorylation.”

**Figure 7.**
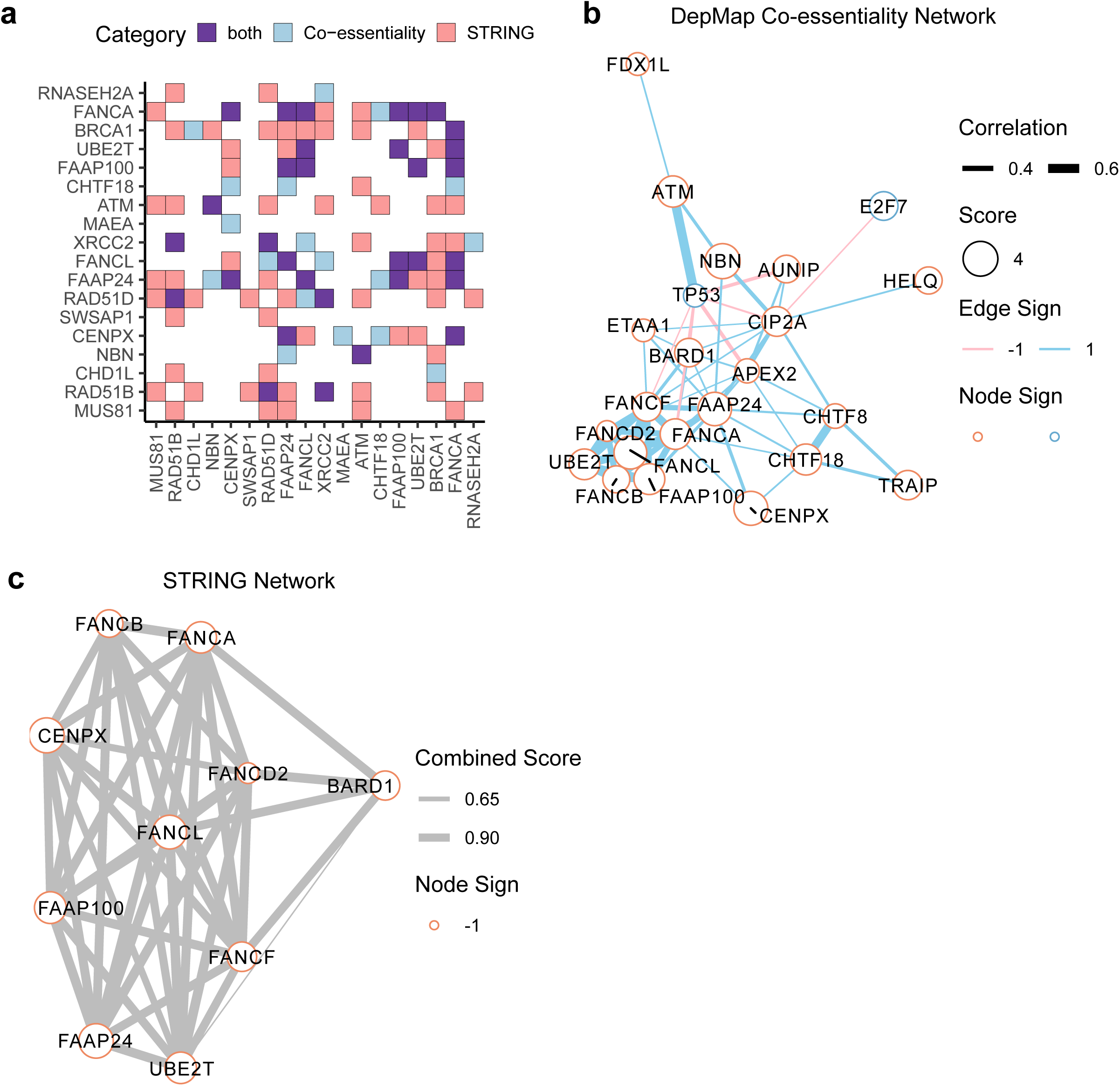
PARP screens reveal functionally coherent clusters of genes. (a) Interactions between the top 20 gene hits from DepMap co-essentiality and STRING network sources. Genes are ordered by absolute average Z-score. (b) Cluster with Fanconi Anemia genes from the DepMap co-essentiality network. Nodes represent genes and the size of each node is proportional to its average Z-score across all screens. Edges represent correlations in DepMap. (c) Cluster with Fanconi Anemia genes in the STRING network. Nodes are the same as (b). Edges represent combined score in STRING.

### Validation

To validate some of these top genes, we performed a competition assay in Meljuso cells in which EGFP labels cells with Cas9 and we assess the fraction of EGFP-positive cells over time by flow cytometry (Supplementary Fig. 8a). We observed that loss of WSB2 or BCL2L1 sensitizes cells to S63845, whereas loss of MARCH5 or MCL1 sensitizes cells to both A-1331852 and navitoclax, another small molecule inhibitor of BCL2L1 (Supplementary Fig. 8b). To further confirm these results, we performed screens using the small molecule inhibitors in a third cell line, the melanoma line A375, using an orthogonal library of Spyo-guides, Gattinara (**Supplementary Data 3**). Gattinara is designed with 2 guides per gene, to reduce the cost of executing these screens, and is complementary to Brunello, in that no targeting guides are shared across these libraries. With the BCL2L1 inhibitor A-1331852 (Fig. 2f), the top 4 sensitizer genes were UBE2J2 (Z-score −13.6), BCL2L2 (−12.7), MCL1 (−10.9), and MARCH5 (−9.8); UBE2K ranked tenth (−6.7). Likewise, with the MCL1 inhibitor S63845 (Supplementary Fig. 9), WSB2 and BCL2L1 scored as the #1 (Z-score −7.7) and #3 (Z-score −6.0) sensitizer hits, respectively, confirming that the strongest genes observed in Meljuso and OVCAR8 cells reproduce in a third cell line with additional guides.

Finally, we attempted an alternative approach to anchor screens, which uses only SpyoCas9 to generate both knockouts (Fig. 4a). Because the two guides are expressed on different vectors but use the same promoters, this system potentially has less competition than approaches that, in order to express two guides on the same vector, use different Pol III promoters^10^. This approach also has the substantial benefit that many Spyo-guides are already well-validated, and only a single Cas9 protein is used. Here, we generated a secondary library targeting 390 potential hit genes that showed evidence of activity in the primary MCL1 and BCL2L1 screens as well as 857 additional non-scoring genes, with 10 guides per gene. We cloned the library into lentiCRISPR-v2 and conducted anchor screens against three Spyo-guides targeting MCL1 in A375 cells, or a control line that was either untreated or treated with S63845. These secondary screens validated BCL2L1 and WSB2 as top synthetic lethal hits with MCL1 (Fig. 4b, c, Supplementary Fig. 10); they also identified three members of a cullin-RING ubiquitin ligase complex, CUL5 (rank 4, Z-score −10.1), RNF7 (rank 3, Z-score −10.5), and UBE2F (rank 5, score −7.6), which are themselves well-correlated in DepMap and were identified as modulators of sensitivity to MCL1 inhibitors in a screen of lung cancer cells in a recent preprint^42^. Only in secondary screens did these three genes score strongly together, highlighting the value of conducting secondary screens with more guides per gene. Another strong hit in the secondary screens was HSP90AB1, a member of the heat shock protein 90 family, which scored strongly in secondary screens with MCL1 knockout (rank 4, Z-score −9.3 across all guides) but did not score with small molecule inhibitors (Gattinara library S63845 rank 1038, Z-score 0.9; secondary library S63845 rank 559, Z-score 0.0). Although it is mechanistically unclear why loss of HSP90 would differentially interact with MCL1 knockout and inhibition, HSP90 has been plausibly linked to both MCL1 and other top hits: HSP90 inhibition results in transcriptional downregulation of MCL1^43^ and CUL5 has been shown to degrade chaperoned proteins following HSP90 inhibition^44^.

Overall, the anchor screens for genetic interactions with BCL2L1 and MCL1 identified both expected and novel partners, and these results were supported by parallel small molecule screens. Further, examination of orthogonal data sources, the STRING and DepMap co-essentiality networks, provides additional confidence for the relevance of these novel interactions and the validity of this approach. We additionally demonstrate that a SpyoCas9-only anchor screening approach can effectively identify synthetic lethal hits and may be a preferable approach for researchers who have already validated effective Spyo-guides targeting their gene of interest.

### Anchor screens with PARP1 knockout and PARP inhibitors

To confirm that this approach can be informative when targeting a gene involved in a different cellular function, we performed anchor screens against the DNA damage response gene PARP1. The synthetic lethal interaction between PARP1 and BRCA1 is well-described^45^, and PARP inhibitors are clinically approved for treatment of tumors with BRCA mutations^46^.

Identifying additional genetic lesions that also synergize with PARP inhibition would thus be valuable, and could expand the population of patients who may benefit from this therapy. We designed two different guides against PARP1 and conducted anchor screens in OVCAR8 and A375 cells; we also performed screens in control cells with the small molecule PARP inhibitor olaparib. Additionally, we acquired and screened a knockout single cell clone of PARP1 in the near-haploid cell line HAP1^47^, as well as a HAP1 parental control line with another PARP inhibitor, talazoparib. Following sequencing, genetic interactions and residual values were calculated as before (**Supplementary Data 1**, **Supplementary Data 2, Table 1**).

Examining top scoring genes across all cell lines and perturbation types (Fig. 5a), we observed that BRCA1 scored as the 18th ranked gene for synthetic lethality with PARP1 (Z-score −3.0), and BRCA2 ranked 54th (Z score −2.4). Conversely, PARG, which catabolizes poly(ADP-ribose), scored as a top buffering gene (rank 2, Z-score 2.8), as has been observed previously^48^. To broadly assess these genetic and small molecule screens, we used several benchmark gene sets (**Supplementary Data 4**): a curated set of genes involved in homologous recombination provided by the Wood laboratory (n = 21)^49^; the Reactome “DNA Repair” gene set (n = 106)^50^; known protein-protein interactors with PARP1 curated by BioGrid (n = 289)^51^; and a high-confidence set of hits identified by Zimmermann and colleagues via CRISPR screens for olaparib sensitivity (n = 73)^52^. Interestingly, the only gene shared in common across these 4 sets is BRCA1 (Fig. 5b). Across all cell lines and perturbation types (Fig. 5c), we saw statistically significant enrichment for each gene set, with Biogrid showing the least (KS statistic 0.21, p-value 9.6×10^-12^) and the Wood lab curated set showing the most (KS 0.80, p-value 1.1×10^-10^; note that p-value calculation is set-size dependent, and the KS statistic is more appropropriate for comparing across gene sets of different sizes).

Comparing genetic knockout of PARP1 to small molecule inhibition, we generally observed concordance (Pearson R = 0.52) but there were outliers (Fig. 6a). For example, XRCC1, which is known to interact directly with PARP1^53^, scored as a top sensitizer for small molecule PARP inhibition (ranked 4, Z-score −5.6), but is a top buffering gene upon genetic knockout (rank 2, Z-score 2.9). Additionally, POLB, which interacts directly with XRCC1^54^, and HPF1, which regulates the activity of PARP1 at serine residues^55, 56^, score as top sensitizers for PARP inhibitors (rank 2, Z-score −6.2 and rank 6, Z-score −5.2 respectively) but do not show any interaction in the genetic knockouts (Z-scores 0.1 and −0.3, respectively). These differences may reflect the distinction between PARP1 still being present in the cell but inactive, in the case of the small molecules, compared to the complete loss of PARP1 protein such that it can no longer assemble into complexes. Additionally, PARP2 may compensate for PARP1 in the case of genetic knockout, but is likely also inhibited by the small molecules. These results emphasize that although in many cases small molecule inhibition phenocopies genetic knockout, as with BCL2L1 and MCL1 presented above, exceptions can arise.

We were also interested in differences between the two small molecules used in this study, olaparib and talazoparib, so these were included as independent arms in the screens conducted in A375 cells with the Gattinara library, described above (**Supplementary Data 3**). Overall, the two molecules gave similar results (Pearson R = 0.50), with many of the same top hits (Fig. 6b). For example, ATM, RAD54L, and NBN rank in the top 10 of sensitizers with both small molecules, whereas TP53, PARG, and TP53BP1 score as resistance genes (ranks 1, 2, and 20, respectively, averaging across the two inhibitors). Some differences emerge between the small molecules, however. For example, PARP1 itself scores as a strong resistance gene with talazoparib only (rank 1, Z-score 8.8), as has been described previously^57^, whereas it shows sensitization with olaparib (rank 152, Z-score −3.0). Further, POLB is a strong sensitizer with talazoparib (rank 2, Z-score −10.2) but scores more weakly with olaparib (rank 352, Z-score −2.4). Conversely, loss of HPF1 strongly synergizes with olaparib (rank 4, Z-score −7.4) and has a weaker phenotype with talazoparib (rank 473, Z-score −2.1). These differences in activity may be due to the fact that talazoparib is the stronger PARP-trapping small molecule^58^.

Interestingly, CUL3 scored as a resistance gene for both PARP inhibitors (resistance rank 9, Z-score 4.2), and loss of this gene has previously been shown to cause resistance to vemurafenib, a BRAF inhibitor, in this cell line^59^. However, loss of another potent vemurafenib resistance gene, MED12, confers sensitivity to PARP inhibitors (sensitization rank 9, Z-score −5.9). Transcriptional profiling to understand the different cell states achieved by CUL3 and MED12 knockout, and how that leads to differential response to BRAF and PARP inhibition, could be an interesting model system for understanding resistance mechanisms more generally^60^.

We also observed cell-line-specific differences. For example, the inosine triphosphatase ITPA is the top-ranked sensitizer in HAP1 cells (Z-score −9.6) but does not score in A375 (Z-score 0.1) or OVCAR8 (−0.5); this gene has been implicated in DNA damage previously, as ITPA normally prevents base analogs from contributing to the pool of free nucleotides, whereas repair following their incorporation leads to DNA single-strand breaks^61^. Indeed, RNAse H2 enzymes, which act to remove ribonucleotides incorporated into DNA, also score strongly in our screens across all cell lines, and their loss has previously been shown to sensitize cells to PARP inhibitors^52^.

Conversely, the DNA ligase LIG1, which repairs breaks in DNA during replication, scores strongly in OVCAR8 (rank 4, Z-score −5.1) but not in A375 (rank 1,420, Z-score −1.1) or HAP1 (rank 17,220, Z-score 1.24). Also in OVCAR8, loss of numerous mitochondrial complex I NDUF genes caused sensitivity to olaparib. Indeed, of the 80 genes in the GO gene set “oxidative phosphorylation,” 24 score in the top 100 (Z-scores < −5.2), whereas this gene set does not show evidence of activity in any other condition (Fig. 6c), including PARP1 knockout in OVCAR8. Thus, both cell context and mode of inhibition may lead to divergent phenotypes that will require additional investigation to understand their different mechanisms. The Shieldin complex (C20orf192, also known as SHLD1; FAM35A, also known as SHLD2; and SHLD3) has recently been characterized by several groups^62, 63^, and loss of these genes has been implicated in resistance to PARP inhibitors. These genes did not score in the screens described here, but notably, those previous screens were performed in BRCA1-deficient cells. Further, it was shown that loss of Shieldin genes did not confer resistance to PARP inhibitors in BRCA-proficient cells^63^. Likewise, DYNLL1, a TP53BP1-interacting protein, has been shown to mediate resistance to PARP inhibitors, but this was only studied in the context of BRCA1-deficient cells^64, 65^.

Of the top 40 synthetic lethal genes, 16 did not appear in any of the four examined gene sets (**Supplementary Data 4**). Some of these may be false positives, or might be captured by other sources of gene sets with plausible relationships to PARP biology. For example, SWSAP1 and ZSWIM7 (ranks 6 and 23 overall, respectively) form a complex and are lesser known Rad51 paralogues. These appear to be required for efficient homologous recombination, and have been indicated especially in meiotic recombination^66–68^. Likewise, the genes CHTF18 and RFC1 (ranks 14 and 40, respectively), are each core members of distinct replication fork complexes that load the DNA polymerase processivity factor, PCNA^69^. Some other top hits, however, have no obvious connection to DNA damage in the literature, such as MAEA (rank 12), a RING-domain containing protein that is part of an E3 ligase complex^70^. This gene is highly correlated in co-essentiality with three genes in the DepMap, UBE2H (R = 0.77), WDR26 (R = 0.71), and YPEL5 (R = 0.70), which rank 558, 86, and 212 in the PARP screens, providing confidence that this novel hit generalizes across cell lines.

Finally, we organized the detected genetic interactions by constructing DepMap and STRING networks (Supplementary Figs. 11, 12). In both cases, there was more connectivity than expected by chance (Supplementary Fig. 13) with substantial interconnectivity of top hits in both networks (Fig. 7a). Both the DepMap (Fig. 7b) and STRING (Fig. 7c) networks have a cluster containing the many Fanconi Anemia genes that score as hits in these screens, including UBE2T, which was recently validated in patients as a causal gene for Fanconi Anemia^71^, as well as TRAIP, which has recently been mechanistically connected to the pathway^72^. The resulting networks illustrate how such network approaches can be useful for suggesting function to less characterized genes.

## DISCUSSION

Here we present a facile approach for generating genome-wide genetic interaction maps for individual genes in cell types of interest using CRISPR technology. By timing the delivery of the anchor perturbation, this approach eliminates the need for single cell cloning, which is typically a major bottleneck for experiments with isogenic cell lines.

Importantly, this approach does not generate true isogenic pairs, as DNA double-strand breaks result in a spectrum of indels^73^. Yet, it is also the case that bystander mutations private to any single cell clone are numerous and thus pairs generated by traditional approaches are also not truly isogenic^74^. The success of this approach rests heavily on the activity of the anchor guide, and thus its performance should be validated before beginning such a screening campaign to find, for example, guides that yield a very high fraction of out-of-frame indels. Further, for a given gene of interest, an effective screening strategy may be to perform fewer replicates with any one guide, but with more unique anchor guides, to mitigate potential off-target effects of a particular guide sequence.

That co-essentiality data from large-scale genetic screening projects^34, 35, 39^ can be used to generate genetic interaction maps has been demonstrated by several groups^36–38^, and these data undoubtedly represent a powerful resource for the scientific community. However, these large-scale screening projects are the result of many dollars and years of effort, and it is not trivial for individual researchers with, for example, a patient-derived cell line or an organoid model, to feed into these pipelines. Further, despite the impressive size of these resources, many tumor types and specific genetic lesions are still poorly represented. Thus, the two-pronged approach described here -- perform an anchor screen, then cluster the hits using co-essentiality data -- enables researchers to uncover genetic interactions with a gene of interest in a biologically relevant cell type, but still leverage the data from these large-scale maps to interpret and prioritize the resulting hit genes.

We expect that the anchor screening approach described here will be useful to understand the genetic landscape of a target before a lead candidate small molecule is identified, and to understand the differences between small molecule targeting of a gene and genetic loss-of-function alleles. One use-case for such screens is to identify biomarkers that may indicate sensitivity to a small molecule. For example, as annotated in the cBioPortal^75^, BCL2L1 and MCL1 are each often amplified in tumors, providing justification for the development of clinical inhibitors (Supplementary Fig. 14). We queried other top synthetic lethal hits, and observed that MARCH5 and UBE2J2 are characterized as deep deletions in a fraction of tumors (Supplementary Fig. 15). Likewise, PARP inhibitors have shown clinical efficacy in tumors with BRCA1 or BRCA2 mutations^46^, which was recapitulated in these screens, and their use has been extended to patients with ATM mutations^76^, a gene that also scored here. Several other genes that scored highly as synthetic lethal in these screens are often categorized as deep deletions across a spectrum of tumor types (Supplementary Fig. 16), and thus it is reasonable to speculate that such tumors may be particularly sensitive to PARP inhibitors. Whether these mutations are important for tumorigenesis or are simply bystander events will require further study, but such genomic features may serve as biomarkers to identify populations that would benefit from treatment with small molecule inhibitors.

Although we have presented only double-knockout screens here, simply altering one or both of the Cas9 proteins should unlock a variety of screening possibilities, as we and others have previously demonstrated in “some-by-some” screens^10, 26^. Anchor screens may be particularly powerful when paired with base editing technologies^77^, as the introduction of defined gene edits via homologous recombination is at least an order of magnitude less-efficient than the generation of knockout alleles, and thus the generation of such isogenic cells has commensurately higher costs. Likewise, anchoring on CRISPRa- or CRISPRi-mediated perturbation of a gene or noncoding regulatory element could shed light on networks of transcriptional regulation. Expanding genetic interaction mapping to include various perturbation types and less tractable cell contexts promises to enhance our capacity to uncover gene function.

## ACKNOWLEDGEMENTS

We thank Amy Goodale, Hans Reuter, Wei Zhen, Briana Fritchman, and Xiaoping Yang for producing guide libraries and lentivirus; Olivia Bare and Yenarae Lee for logistics support; Matthew Greene, Adam Brown, Doug Alan, Mark Tomko, and Tom Green for software engineering support; Federica Piccioni and David Root for helpful conversations; the Broad Institute Genomics Platform Walk-up Sequencing group for Illumina sequencing; and the Functional Genomics Consortium for funding support.

## AUTHOR CONTRIBUTIONS

Conceived of the study: JGD

Executed genetic screens: KRS, REH, AKS, CS

Performed analyses: JGD, PCD, MH, NSP

Created visualizations: PCD, KRS

Curated data: PCD, MH

Wrote the manuscript: JGD, PCD, REH (with input from all authors)

Supervised the project: JGD

## DATA AVAILABILITY

The read counts for all screening data and subsequent analyses are provided as Supplementary Data.

## CODE AVAILABILITY

All custom code used for analysis and example notebooks are available on GitHub: https://github.com/PeterDeWeirdt

## METHODS

### Vectors

Individual sgRNA sequences are provided in **Supplementary Table 1**. The following vectors were used in the study and will be made available on Addgene:

pXPR_213 (anchor vector): H1 promoter expresses customizable Saur-guide; EF1a promoter expresses SpyoCas9 and 2A site provides blasticidin resistance (Addgene # TBD).

pXPR_212 (library vector): U6 promoter expresses customizable Spyo-guide; EFS promoter expresses SaurCas9 and 2A site provides puromycin resistance (Addgene # TBD).

pRDA_186 (Spyo-only anchor vector): U6 promoter expresses customizable Spyo-guide; PGK promoter expresses blasticidin resistance and 2A site provides EGFP (Addgene # TBD).

lentiCRISPRv2 (pXPR_023): EF1a promoter expresses SpyoCas9 and 2A site provides puromycin resistance; U6 promoter expresses customizable Spyo-guide (Addgene # 52961).

pRosetta_v2: PGK promoter expresses hygromycin resistance, T2A site provides blasticidin resistance, P2A site provides puromycin resistance, and F2A site provides EGFP (Addgene # 59700).

pLX_311-Cas9: SV40 promoter expresses blasticidin resistance; EF1a promoter expresses SpyoCas9 (Addgene # 96924).

pRDA_118 (modified lentiGuide): U6 promoter expresses customizable Spyo-guide; EF1a promoter provides puromycin resistance (Addgene # TBD). This vector is a derivative of the lentiGuide vector, with minor modifications to the tracrRNA.

pRDA_103: H1 promoter with two Tet operator (TetO) sites expresses customizable Spyo-guide; short EF1a promoter (EFS) expresses SaurCas9, 2A provides TetR, and 2A provides blasticidin resistance (Addgene # TBD).

pXPR_124: EF1a promoter expresses SpyoCas9 and P2A provides EGFP (Addgene # TBD).

### Library production

Library production was performed as described previously^28^.

### Lentivirus production

Lentivirus production was performed as described previously^28^.

### Cell culture

All cell lines were maintained in humidity-controlled, 37°C incubators with 5.0% CO_2_ and were passaged every 2-4 days. Cell lines were routinely tested for and found to be free of mycoplasma contamination via a PCR assay. A375, Meljuso, and OVCAR8 cell lines were obtained from the Cancer Cell Line Encyclopedia. HEK293T cells were obtained from ATCC (CRL-3216) many years ago, and population expanded that happened to adhere better to plasticware. HAP1 parental and PARP1-knockout cells were obtained from Horizon Discovery.

Cells were regularly maintained in antibiotic-free media, except during screens, when cells were maintained in media containing 1% penicillin/streptomycin. The following media conditions and doses of polybrene, puromycin, and blasticidin, respectively, were used:

A375: RPMI + 10% FBS; 1 µg/mL; 1 µg/mL; 5 µg/mL

HAP1: IMDM + 10% FBS; 4 µg/mL; 1 µg/mL; 5 µg/mL

HEK293T: DMEM + 10% FBS; N/A; N/A; N/A

Meljuso: RPMI + 10% FBS; 4 µg/mL; 1 µg/mL; 4 µg/mL

OVCAR8: RPMI + 10% FBS; 4 µg/mL; 1 µg/mL; 8 µg/mL

Olaparib (10621) was obtained from Cayman Chemical Co. Talazoparib (BMN 673), navitoclax (ABT-263), and venetoclax (ABT-199) were obtained from Selleckchem. S63845 was a gift from Guo Wei. A-1331852 (A-6048) was obtained from Active Biochem.

### Determination of antibiotic dose

In order to determine an appropriate antibiotic dose for each cell line, cells were transduced with the pRosetta_v2 lentivirus such that approximately 30% of cells were infected and therefore EGFP+. At least 1 day post-transduction, cells were seeded into 6-well dishes at a range of antibiotic doses (e.g. from 0 µg/mL to 8 µg/mL of puromycin). The rate of antibiotic selection at each dose was then monitored by performing flow cytometry for EGFP+ cells. For each cell line, the antibiotic dose was chosen to be the lowest dose that led to at least 95% EGFP+ cells after antibiotic treatment for 7 days (for puromycin) or 14 days (for blasticidin and hygromycin).

### Determination of lentiviral titer

To determine lentiviral titer for transductions, cell lines were transduced in 12-well plates with a range of virus volumes (e.g. 0, 150, 300, 500, and 800 μL virus) with 3.0 × 10^6^ cells per well in the presence of polybrene. The plates were centrifuged at 640 x g for 2 h and were then transferred to a 37 °C incubator for 4–6 h. Each well was then trypsinized, and an equal number of cells seeded into each of two wells of a 6-well dish. Two days post-transduction, puromycin was added to one well out of the pair. After 5 days, both wells were counted for viability. A viral dose resulting in 30–50% transduction efficiency, corresponding to an MOI of ∼0.35–0.70, was used for subsequent library screening.

### Screens

Cells expressing the anchor construct were made by transducing with one of the anchor lentiviral vectors and selecting with blasticidin. They were then transduced with library (e.g. Brunello in pXPR_212) in two biological replicates at a low MOI (∼0.5). Transductions were performed with enough cells to achieve a representation of at least 500 cells per sgRNA per replicate, taking into account a 30–50% transduction efficiency. Throughout the screen, cells were split at a density to maintain a representation of at least 500 cells per sgRNA, and cell counts were taken at each passage to monitor growth. Puromycin selection was added 2 days post-transduction and was maintained for 5–7 days. 3 weeks post-transduction, cells were pelleted by centrifugation, resuspended in PBS, and frozen promptly for genomic DNA isolation.

### Genomic DNA isolation and sequencing

Genomic DNA (gDNA) was isolated using the Machery Nagel NucleoSpin Blood Maxi (2e7–1e8 cells), Midi (5e6–2e7 cells), or Mini (<5e6 cells) kits as per the manufacturer’s instructions. The gDNA concentrations were quantitated by Qubit. For PCR amplification, gDNA was divided into 100 μL reactions such that each well had at most 10 μg of gDNA. Per 96 well plate, a master mix consisted of 150 μL ExTaq DNA Polymerase (Takara), 1 mL of 10x Ex Taq buffer, 800 μL of dNTP provided with the enzyme, 50 μL of P5 stagger primer mix (stock at 100 μM concentration), and 2 mL water. Each well consisted of 50 μL gDNA plus water, 40 μL PCR master mix, and 10 μL of a uniquely barcoded P7 primer (stock at 5 μM concentration). For the Spyo-only validation screens in A375 cells, the master mix was modified as follows: 150 μL Titanium Taq DNA Polymerase (Takara), 1 mL of 10x Titanium Taq buffer, 800 μL of dNTP (Takara, 4030), 50 μL of P5 stagger primer mix (stock at 100 μM concentration), 500 μL of DMSO, and 1500 μL water. We recommend the latter protocol going forward.

PCR cycling conditions: an initial 1 min at 95 °C; followed by 30 s at 94 °C, 30 s at 52.5 °C, 30 s at 72 °C, for 28 cycles; and a final 10 min extension at 72 °C. P5/P7 primers were synthesized at Integrated DNA Technologies (IDT). PCR products were purified with Agencourt AMPure XP SPRI beads according to manufacturer’s instructions (Beckman Coulter, A63880). Samples were sequenced on a HiSeq2500 HighOutput (Illumina), loaded with a 5% spike-in of PhiX DNA.

Reads were counted by first searching for the CACCG sequence in the primary read file that appears in the vector 5′ to all sgRNA inserts. The next 20 nts are the sgRNA insert, which was then mapped to a reference file of all possible sgRNAs present in the library. The read was then assigned to a condition (e.g. a well on the PCR plate) on the basis of the 8nt barcode included in the P7 primer.

### Screen analysis

Following deconvolution, the resulting matrix of read counts was first normalized to reads per million within each condition by the following formula: read per sgRNA/total reads per condition × 10^6^. Reads per million was then log2-transformed by first adding one to all values, which is necessary in order to take the log of sgRNAs with zero reads. For each sgRNA, the log2-fold-change from plasmid DNA (pDNA) was then calculated.

The log2-fold-changes for each perturbed arm were fit using a natural cubic spline with three degrees of freedom, using the log2-fold-changes of the relevant control arm as reference. We then used the residual from this fit as a phenotypic measure for each guide.

### Synthetic interaction statistical significance

In order to determine the significance of synthetic interactions at the gene level we used a Z-test. We averaged the residuals of guides targeting a gene and then calculated a Z-score for these values using the average residual and standard deviation of all guides. In doing so, we assume the distribution of residuals is normal and the average and standard deviation of all guides is representative of the population.

### Network analysis

All network analyses were done in R. Visualizations were done using the tidygraph and ggraph packages. Network clustering was done using the cluster_louvain function in igraph^78^. We used absolute correlations for co-essentiality and combined scores for STRING as edge weights for the clustering algorithm. Graphs are plotted using the force directed layout in igraph.

### GFP competition assay

Doxycycline inducible MCL1, BCL2L1, MARCH5 and WSB2 anchor cell lines were generated by delivering the pRDA_103 vector via a no-spin transduction. Meljuso cells were seeded in a T75 flask in a total volume of 8.6 mL of virus-containing media with polybrene at 0.5 μg mL^−1^.

Flasks were then transferred to an incubator overnight, and the virus-containing media was replaced with fresh media 16–18 h after seeding. Blasticidin selection was added 2 days post-transduction and was maintained for 14 days. Cells were then transduced with pXPR_124 (SpyoCas9-P2A-EGFP) at an MOI of ∼0.5 using the spin method, generating a mixed population of EGFP+ and EGFP-cells. After 5 days, cells were treated with 1 μg mL^−1^ of doxycycline to induce expression of Spyo-guide. On day 7 post infection, cells were treated with 250 nM of either S63845, venetoclax (ABT-263), A-1331852, or navitoclax (ABT-199). The fraction of EGFP-positive cells was monitored for 2 weeks by flow cytometry (BD Accuri C6 Sampler) upon every cell passage.

---

**Table 1** Genome-wide screens in this study. B = Brunello library; G = Gattinara library; A133 = A-1331852; Olap. = Olaparib; Talaz. = Talazoparib. Pearson correlation coefficients are shown for the fold-change (log_2_) relative to the plasmid DNA for replicate screens.

**Supplementary Figure 1.**
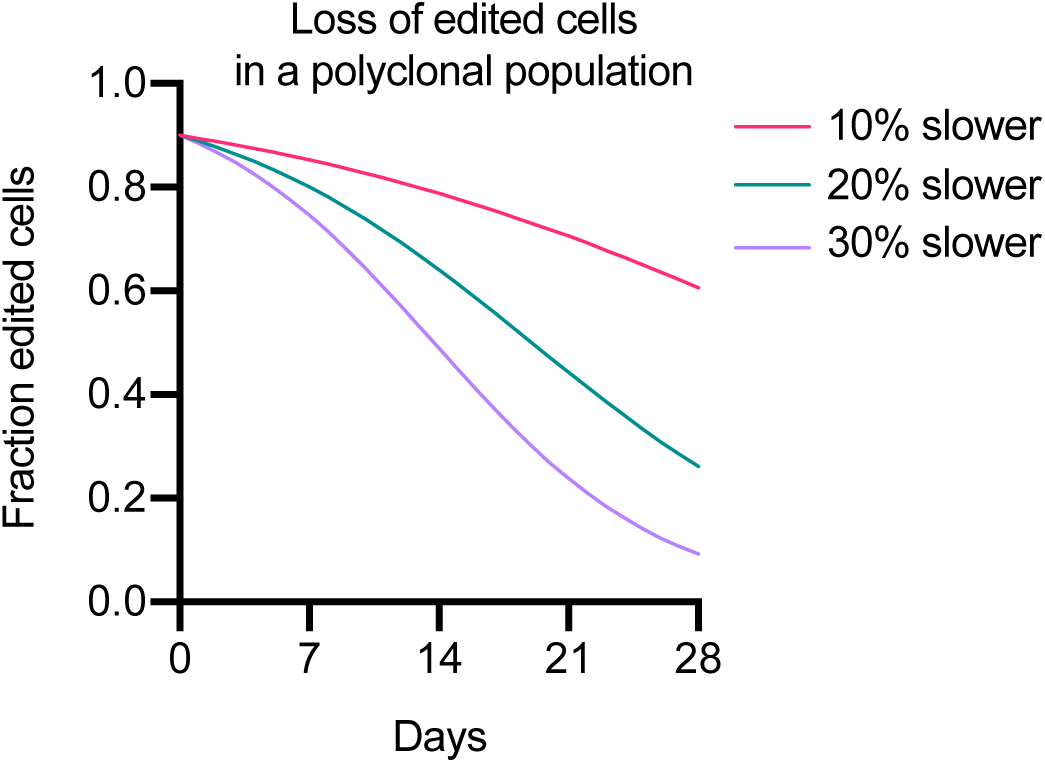
Simulation of competition between edited and unedited cells in a polyclonal population. Edited cells are outcompeted over the span of 28 days assuming the edit causes cells to double 10%, 20%, or 30% more slowly than unmodified cells, and assuming the unedited cells have a doubling time of 24 hours.

**Supplementary Figure 2.**
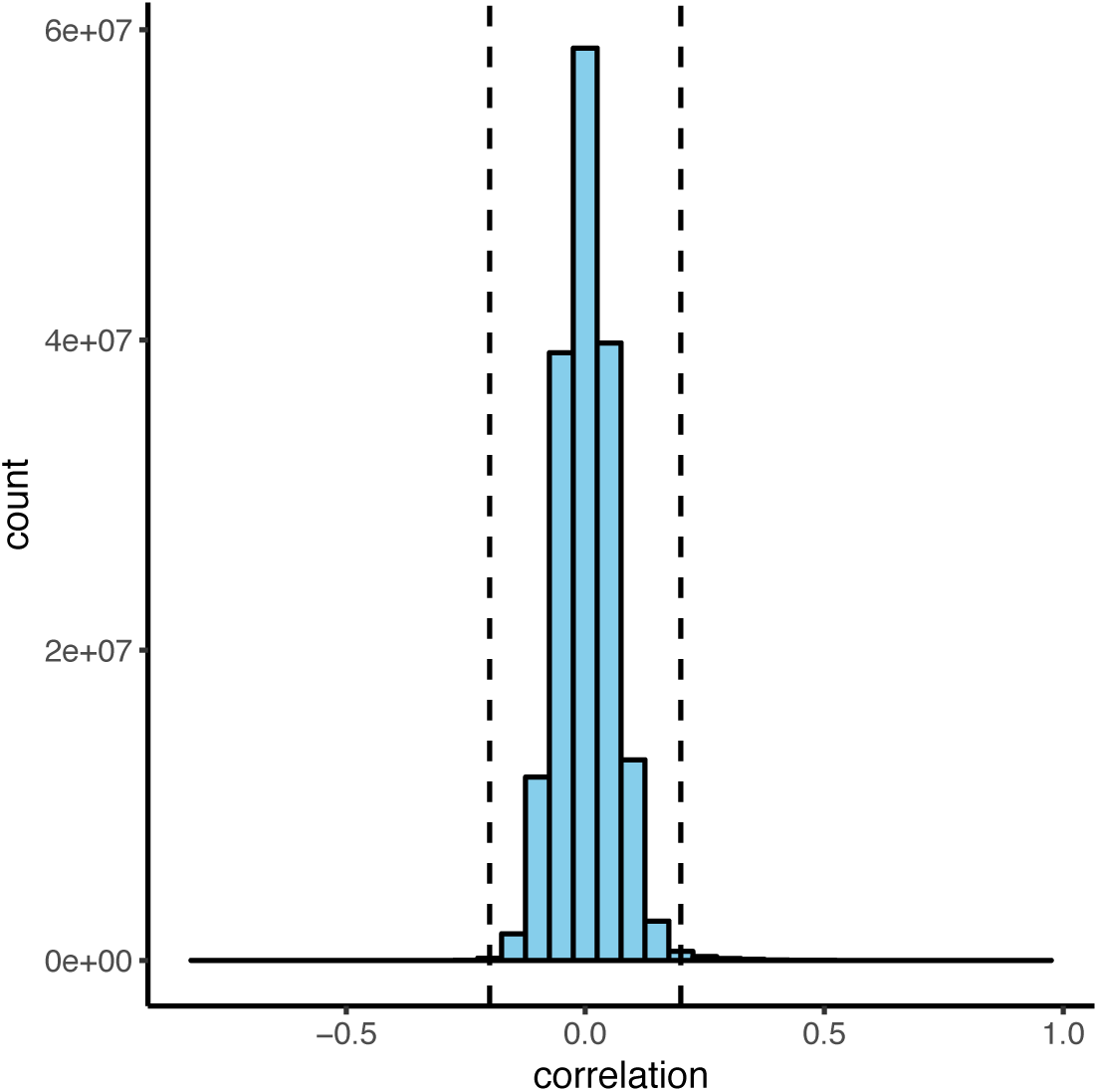
Co-essentiality correlations less than −0.2 or greater than 0.2 are rare in DepMap. Histogram of all correlations using a binwidth of 0.05.

**Supplementary Figure 3.**
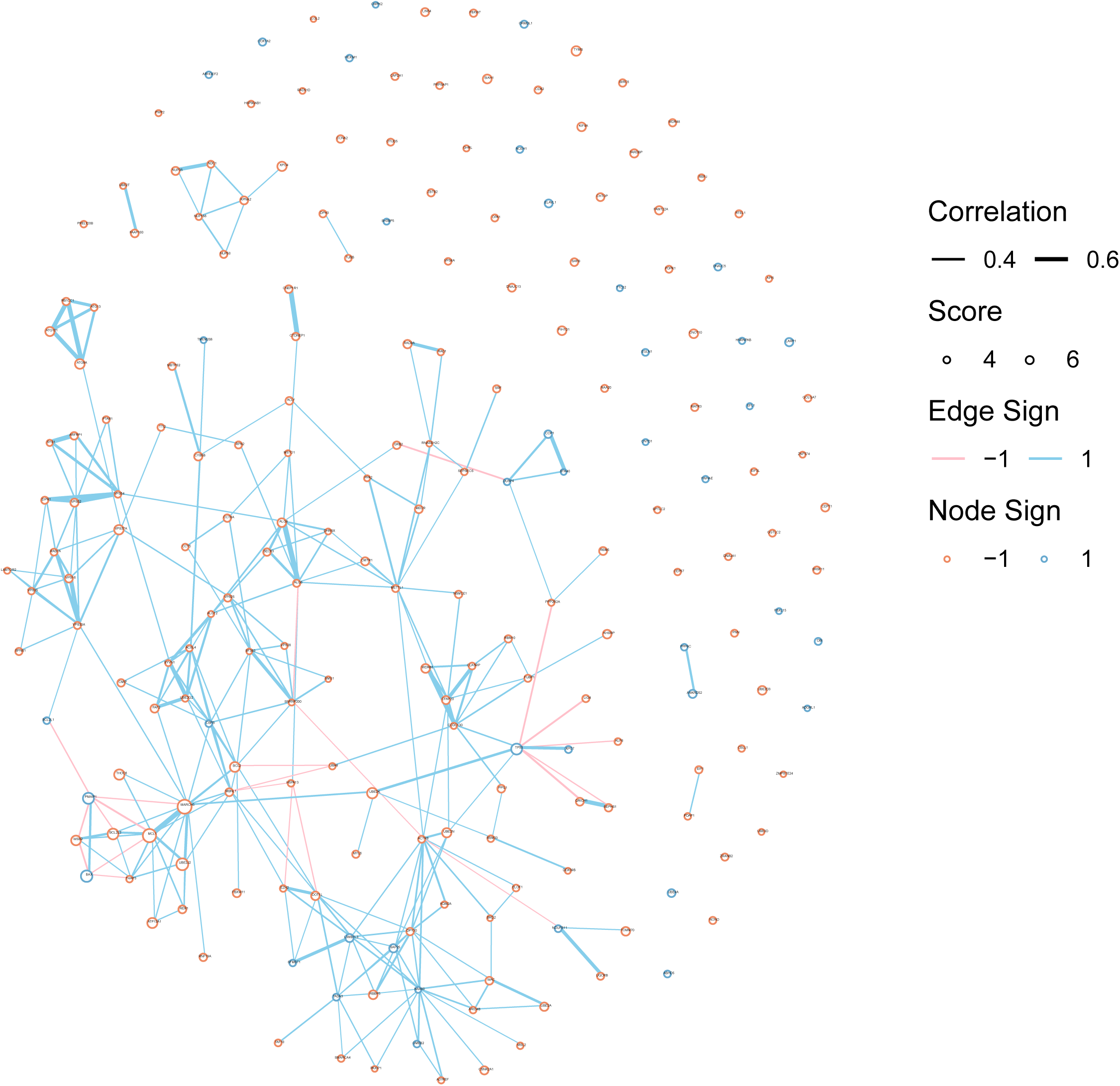
Co-essentiality correlations from DepMap connect functionally related gene hits from BCL2L1 anchor screens. Nodes represent genes and the size of each node is proportional to its average Z-score across all screens. Edges represent correlations in DepMap.

**Supplementary Figure 4.**
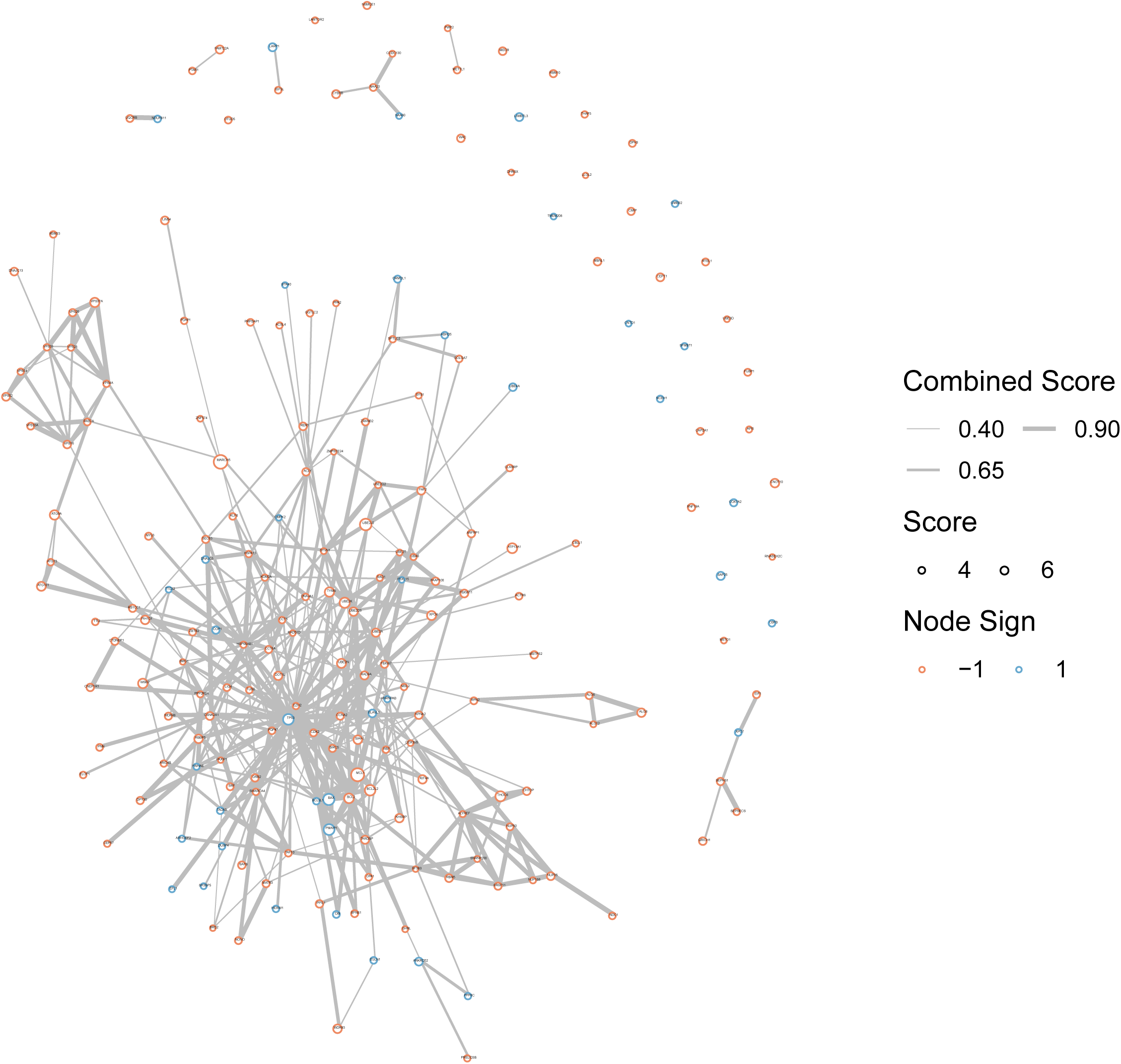
Combined scores from STRING connect functionally related gene hits from BCL2L1 anchor screens. Nodes represent genes and the size of each node is proportional to its average Z-score across all screens. Edges represent combined score in STRING.

**Supplementary Figure 5.**
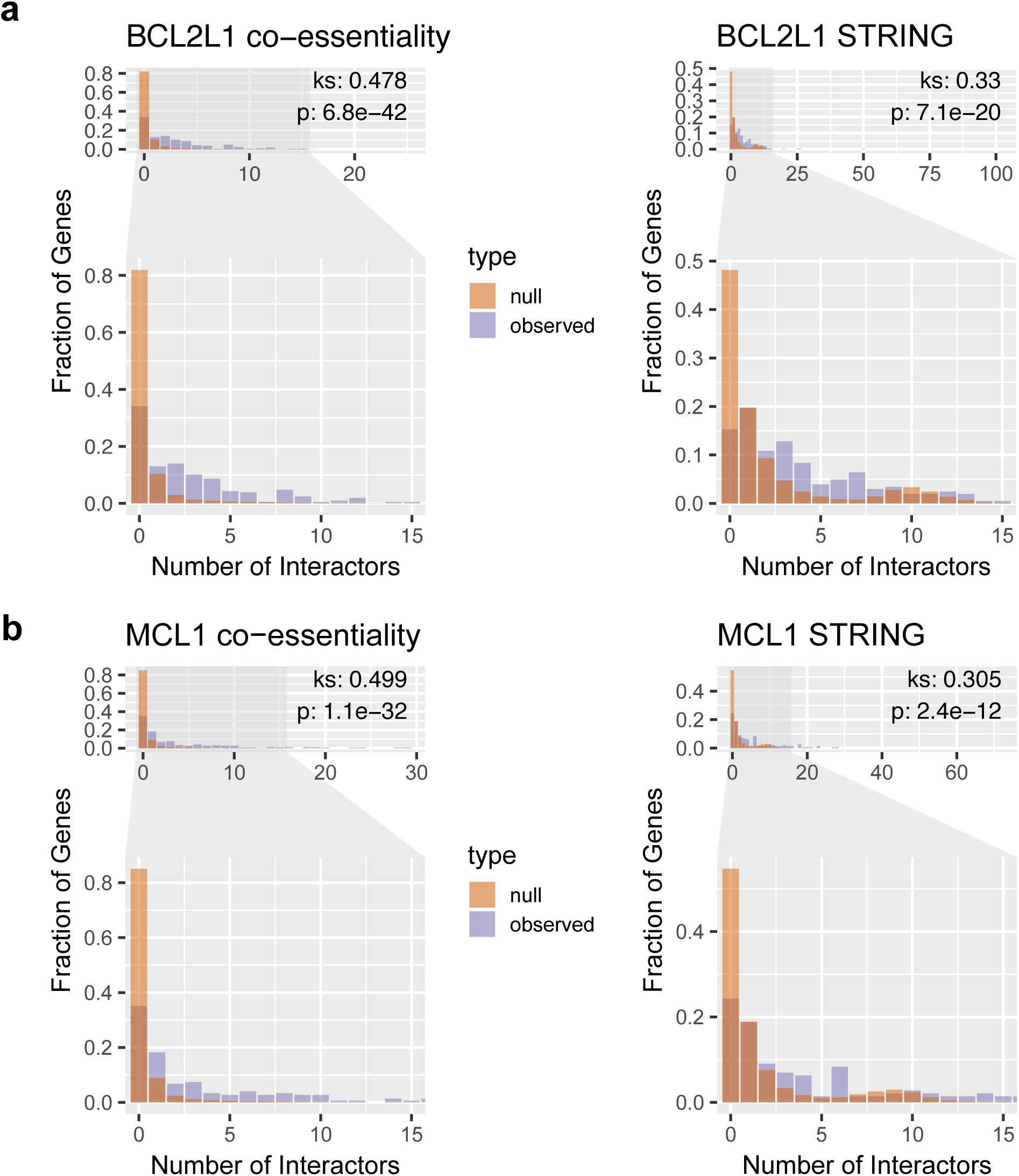
Hits from the BCL2L1 and MCL1 anchor screens are enriched in interactions. (a) Degree distribution for the observed network of BCL2L1 hits compared with a null distribution using DepMap co-essentialities and STRING combined scores. We average 1,000 random networks, each of which has the same number of genes as the original network, to generate the null. To determine statistical significance we used a KS-test with the alternative hypothesis that the observed cumulative distribution was less than the null. (b) Same as (a) but for MCL1.

**Supplementary Figure 6.**
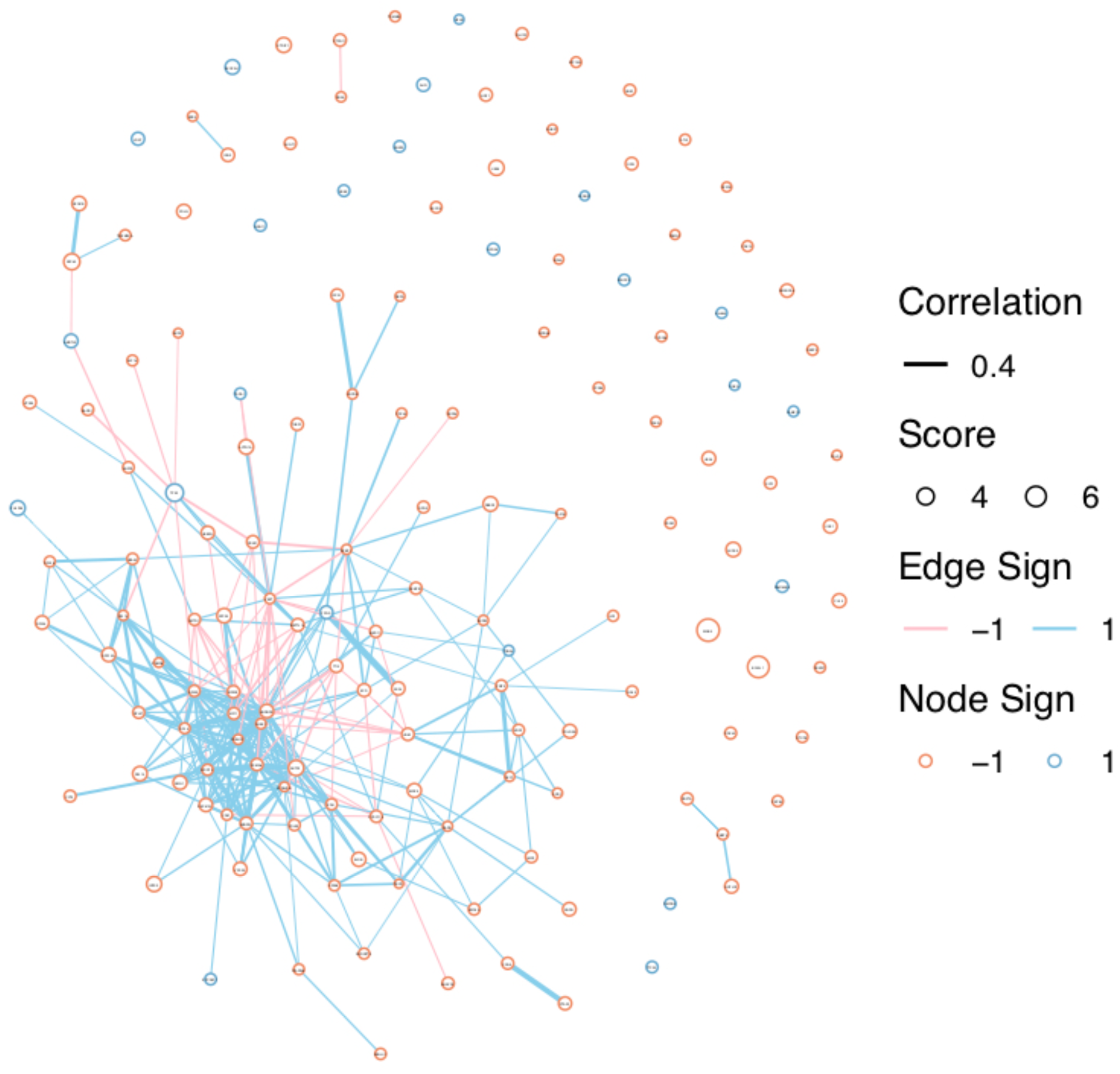
Co-essentiality correlations from DepMap connect functionally related gene hits from MCL1 anchor screens. Nodes represent genes and the size of each node is proportional to its average Z-score across all screens. Edges represent correlations in DepMap.

**Supplementary Figure 7.**
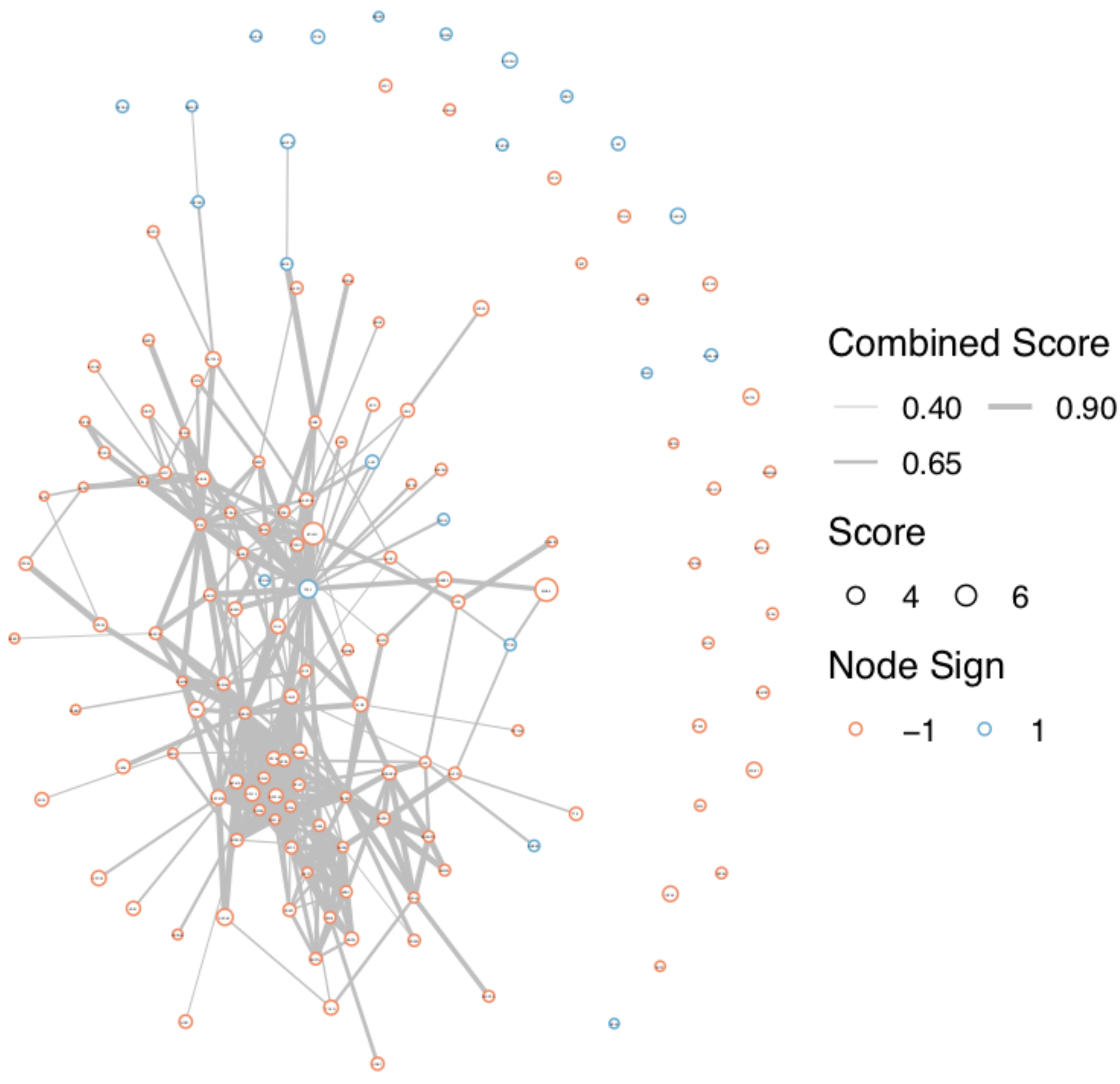
Combined scores from STRING connect functionally related gene hits from MCL1 anchor screens. Nodes represent genes and the size of each node is proportional to its average Z-score across all screens. Edges represent combined score in STRING.

**Supplementary Figure 8.**
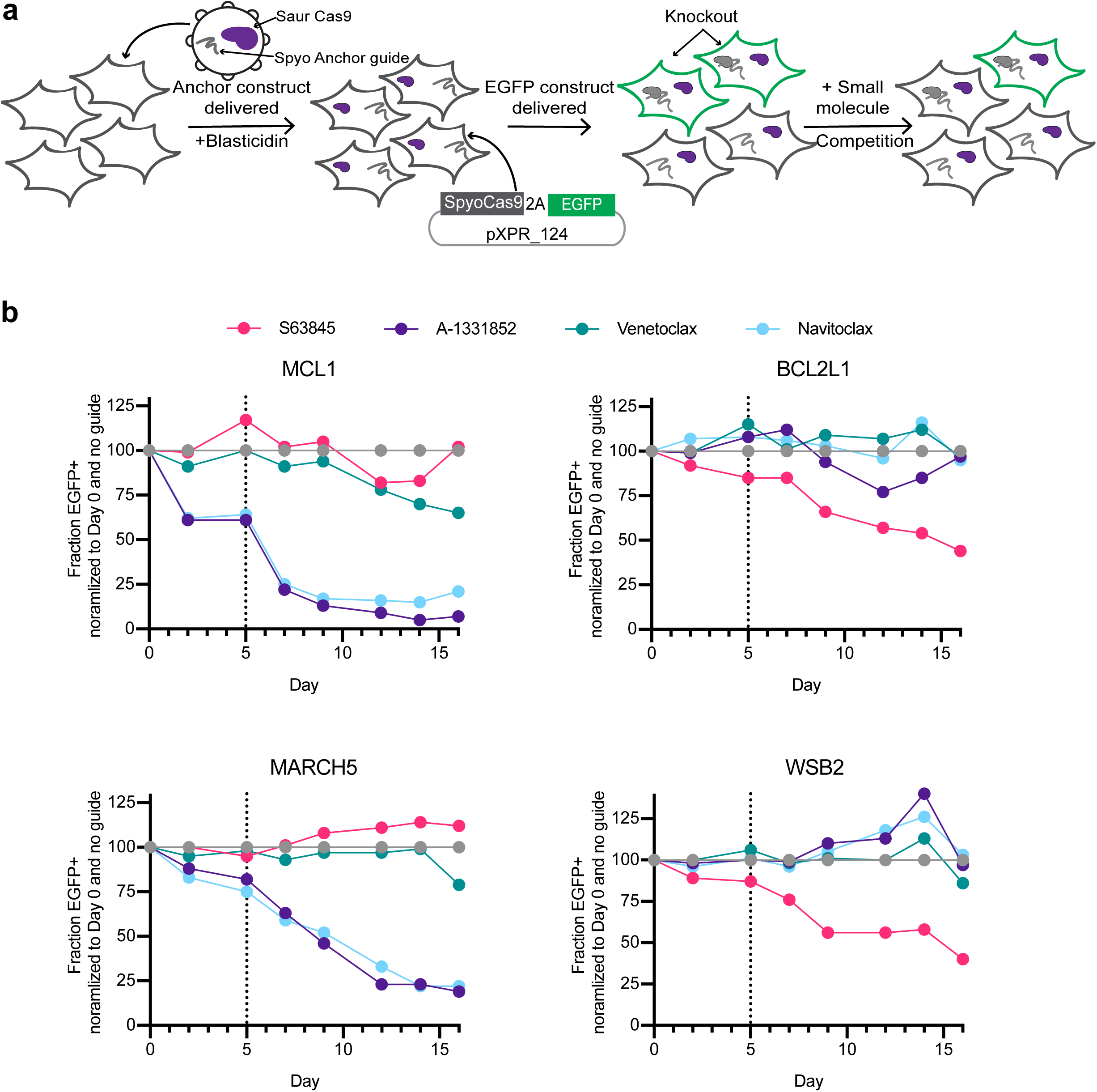
EGFP competition assay validates selected genetic interactions. (a) Schematic of the competition experiment. First, individual Spyo-guides targeting MCL1, WSB2, BCL21, MARCH, or a control guide are delivered in a dox-inducible vector, pRDA_103. Next, Spyo-Cas9 and EGFP are co-delivered via the pXPR_124 vector, and cells are treated with small molecule inhibitors. Although SaurCas9 is delivered with the anchor vector, it is not used in this experimental set-up. (b) Percentage of EGFP-positive cells over time in knockout populations treated with small molecule inhibitors, normalized first to the zero time point for each guide, the time of small molecule addition, and then to the same treatment in the cells infected with a control guide. Doxycycline was added at day 5, indicated by the vertical dotted line.

**Supplementary Figure 9.**
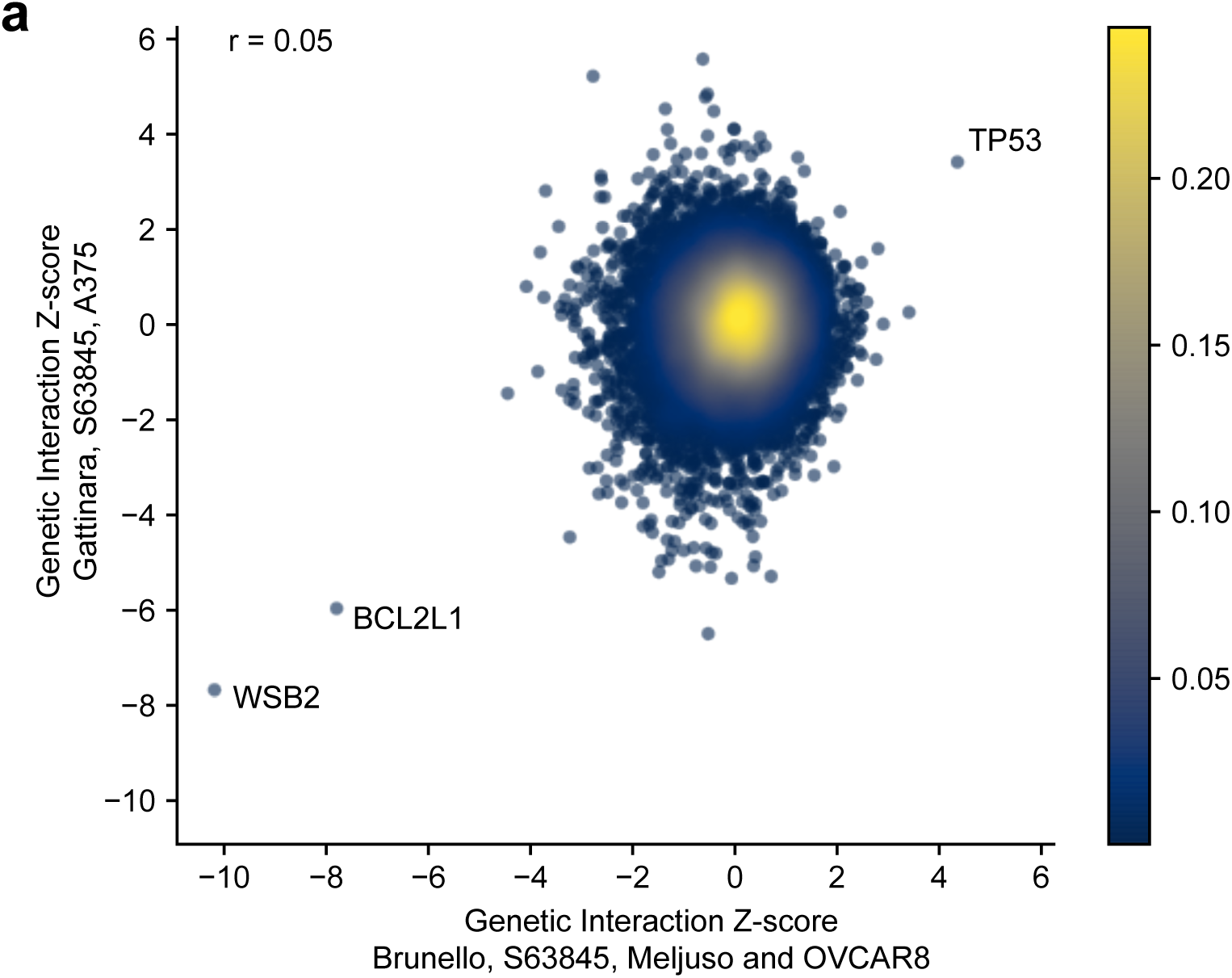
Gattinara and Brunello screened with S63845 identify similar MCL1 genetic interactions. (a) Z-scores for S63845 screened with Brunello and averaged for Meljuso and OVCAR8 cells compared to Z-scores for S63845 screened with Gattinara in A375. Points are colored by density. Pearson correlation coefficient is indicated.

**Supplementary Figure 10.**
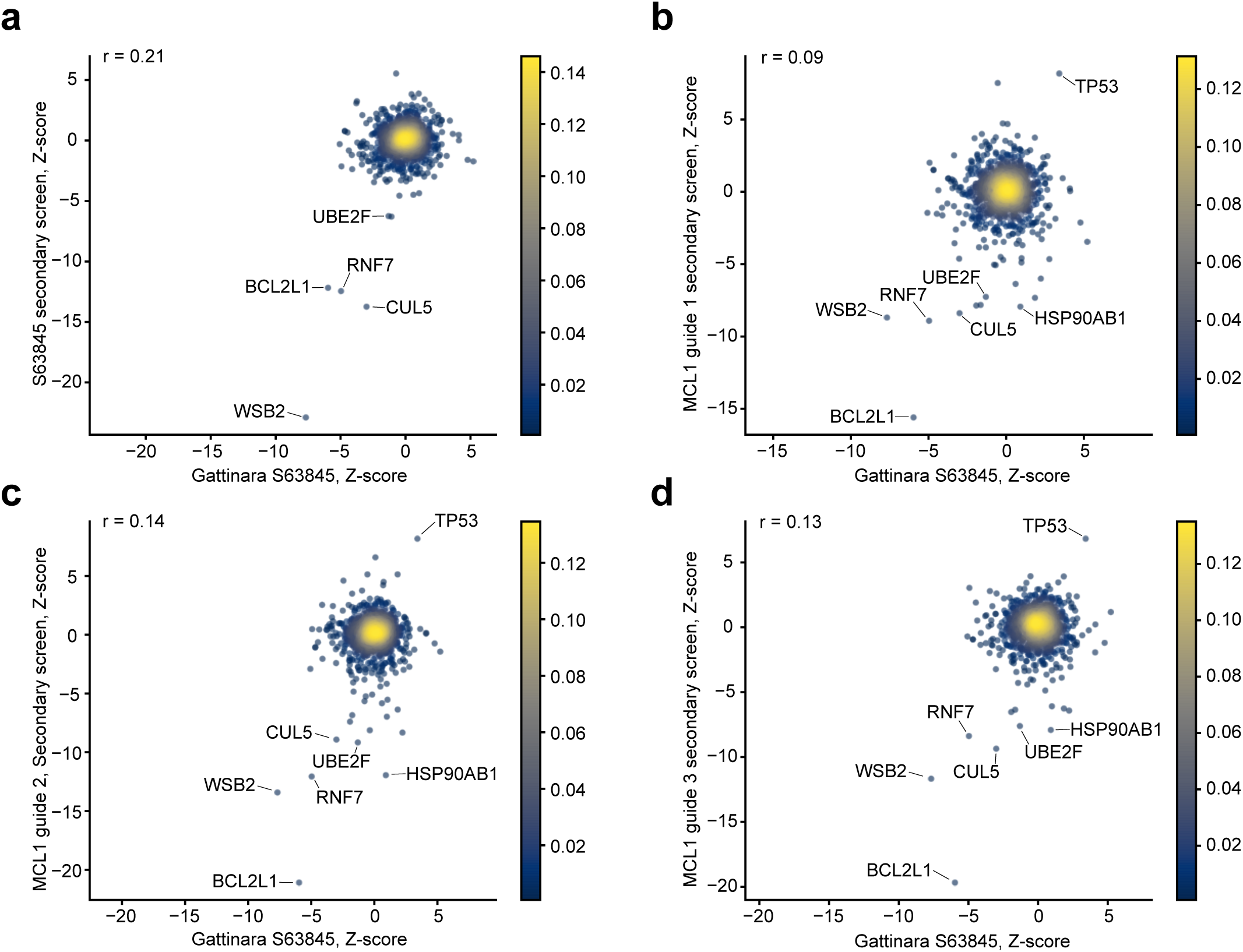
Gattinara and secondary screens identify similar MCL1 genetic interactions. (a) Comparison of Z-scores for S63845 screened with Gattinara and the secondary library. Pearson correlation is reported and points are colored by density. (b) Comparison of Z-scores for S63845 screened with Gattinara and guide 1 of three MCL1 anchor guides screened with the secondary library. Pearson correlation is reported and points are colored by density. (c) Comparison of Z-scores for S63845 screened with Gattinara and guide 2 of three MCL1 anchor guides screened with the secondary library. Pearson correlation is reported and points are colored by density. (d) Comparison of Z-scores for S63845 screened with Gattinara and guide 3 of three MCL1 anchor guides screened with the secondary library. Pearson correlation is reported and points are colored by density.

**Supplementary Figure 11.**
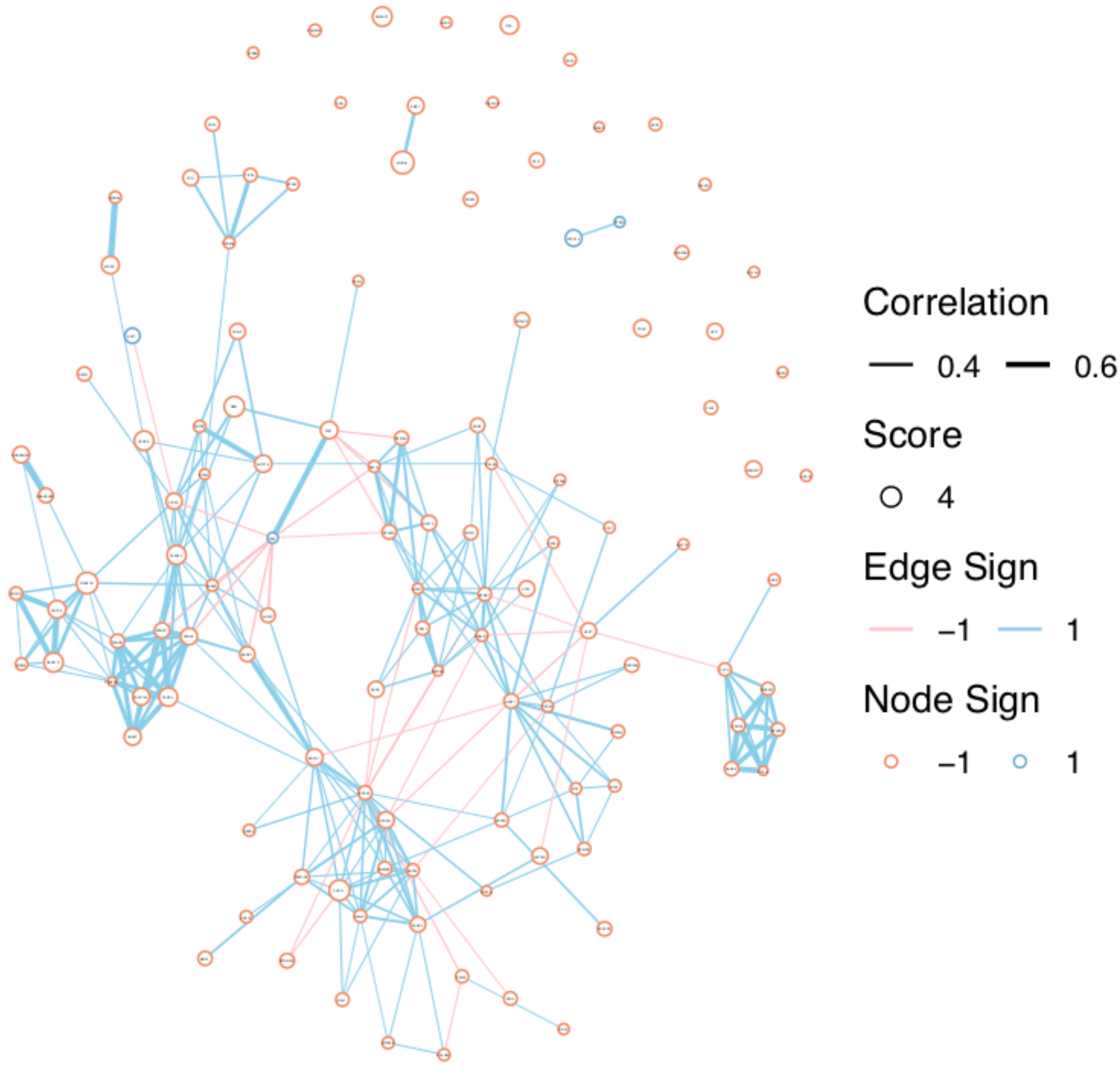
Co-essentiality correlations from DepMap connect functionally related gene hits from PARP anchor screens. Nodes represent genes and the size of each node is proportional to its average Z-score across all screens. Edges represent correlations in DepMap.

**Supplementary Figure 12.**
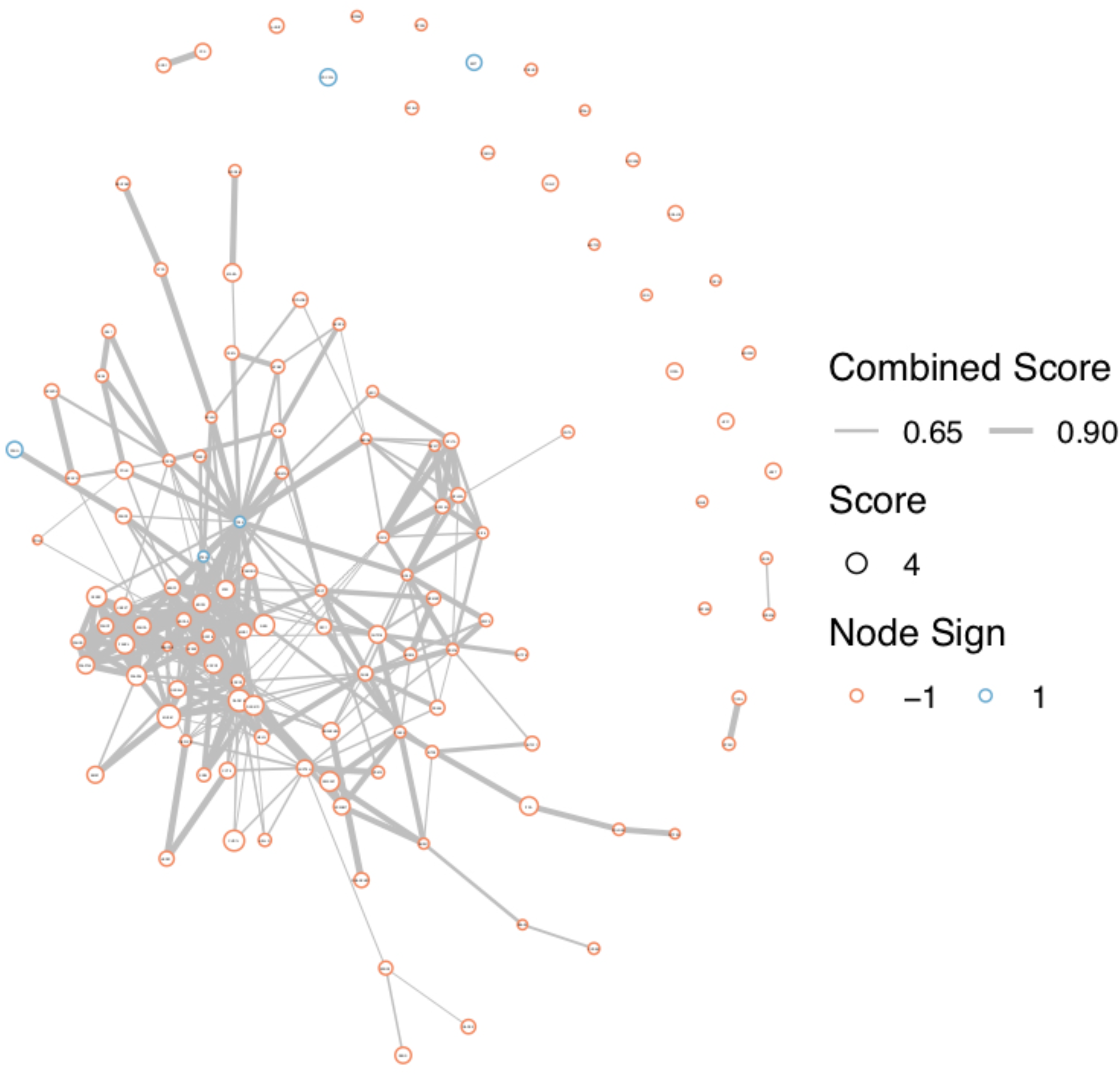
Combined scores from STRING connect connect functionally related gene hits from PARP anchor screens. Nodes represent genes and the size of each node is proportional to its average Z-score across all screens. Edges represent combined score in STRING.

**Supplementary Figure 13.**
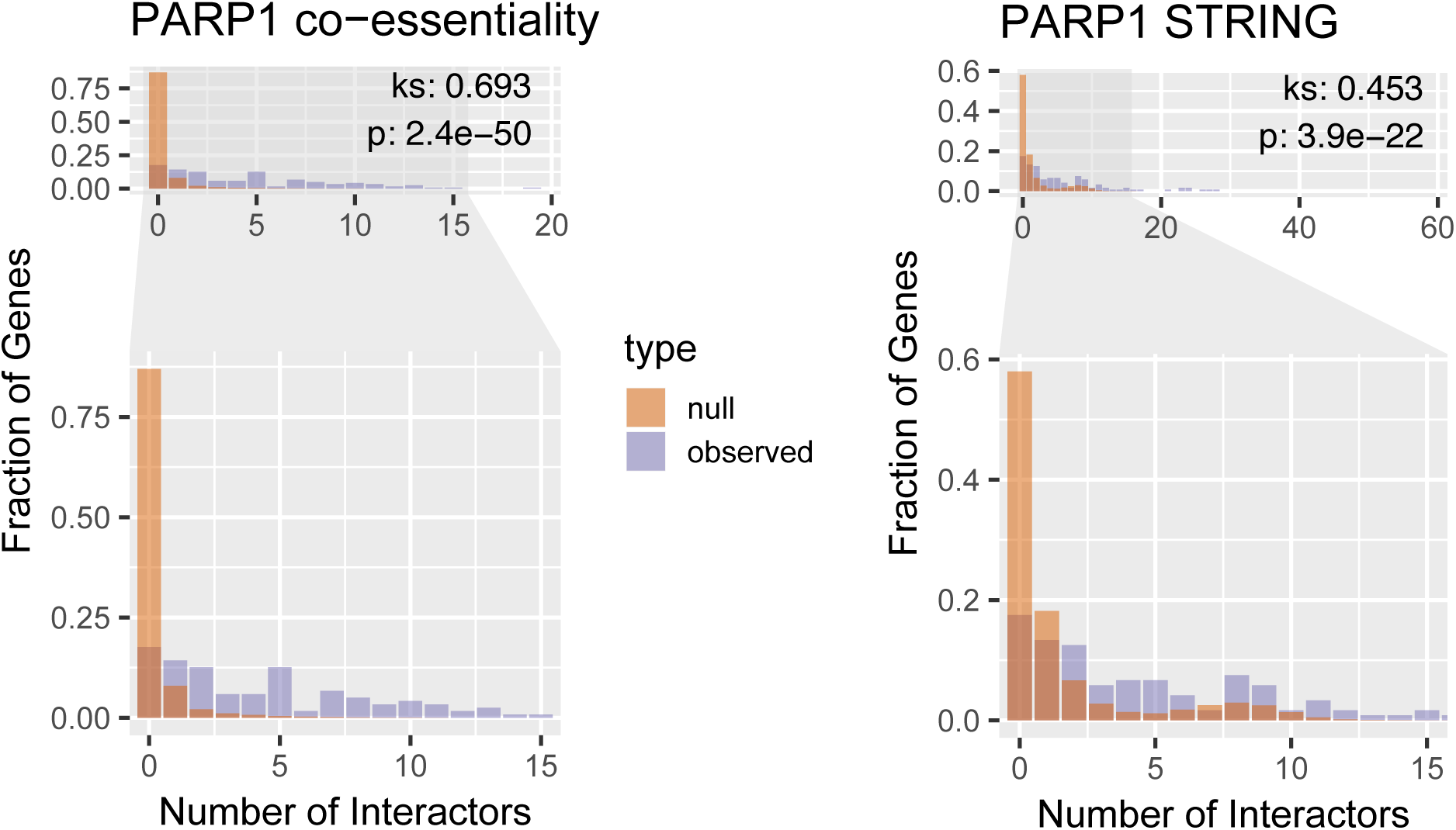
Hits from the PARP anchor screens are enriched in interactions. Degree distribution for the observed network of PARP1 hits compared with a null distribution using DepMap co-essentialities and STRING combined scores. We average 1,000 random networks, each of which has the same number of genes as the original network, to generate the null. To determine statistical significance we used a KS test with the alternative hypothesis that the observed cumulative distribution was less than the null.

**Supplementary Figure 14.**
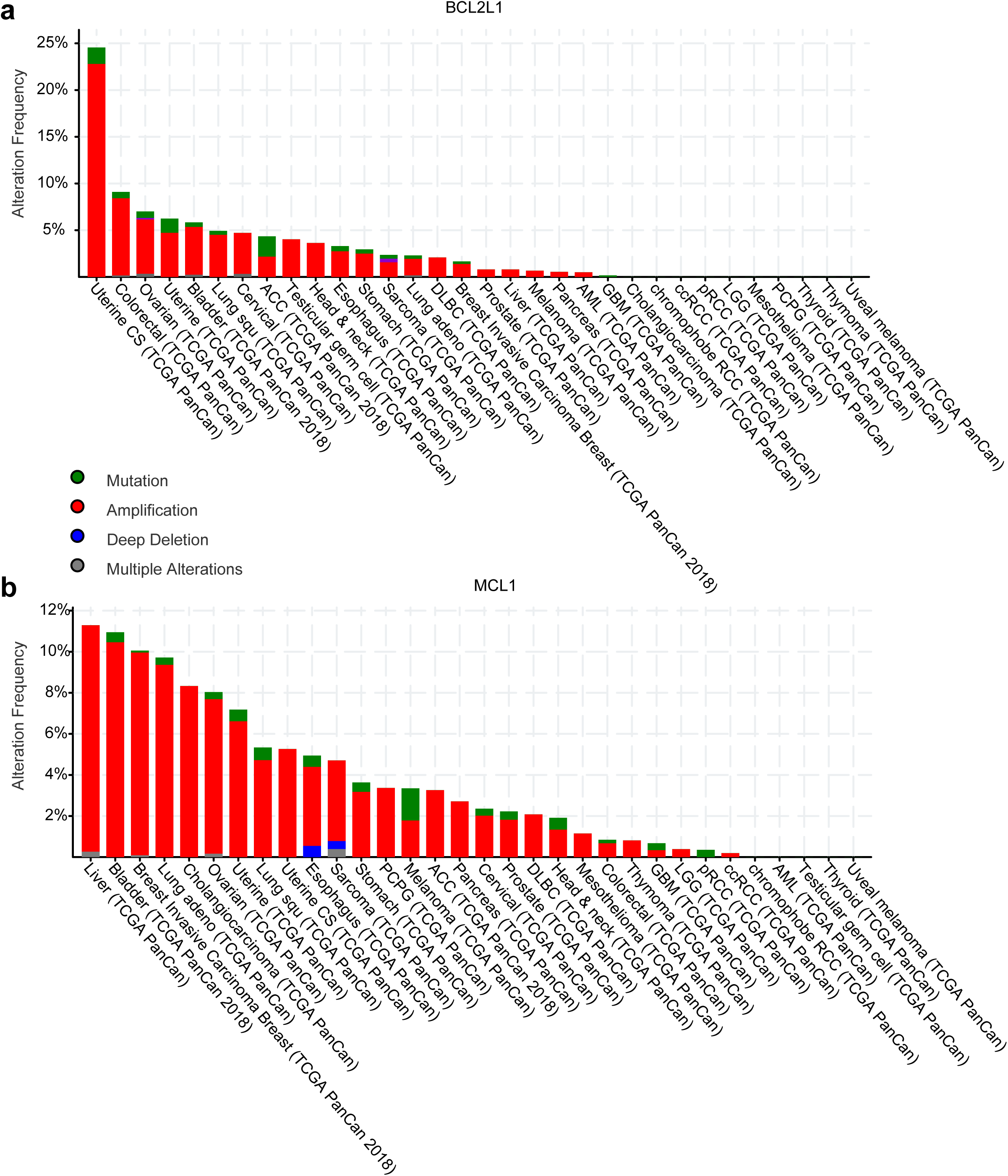
Alteration Frequency plots from cBioPortal for BCL2L1 and MCL1. The TCGA PanCancer Atlas Studies were queried via the web interface for these genes, and the resulting plots are shown here.

**Supplementary Figure 15.**
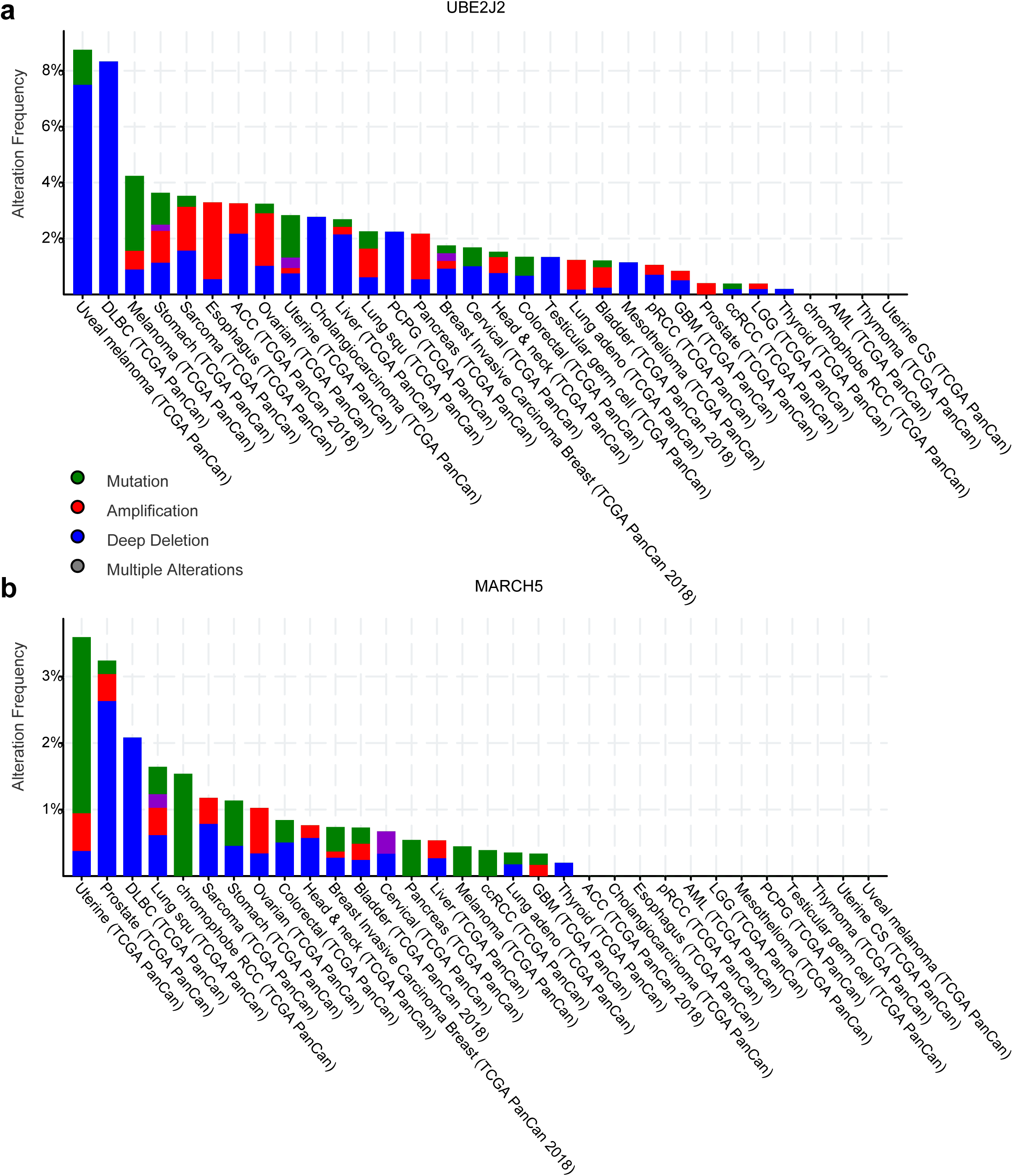
Alteration Frequency plots from cBioPortal for UBE2J2 and MARCH5. The TCGA PanCancer Atlas Studies were queried via the web interface for these genes, and the resulting plots are shown here.

**Supplementary Figure 16.**
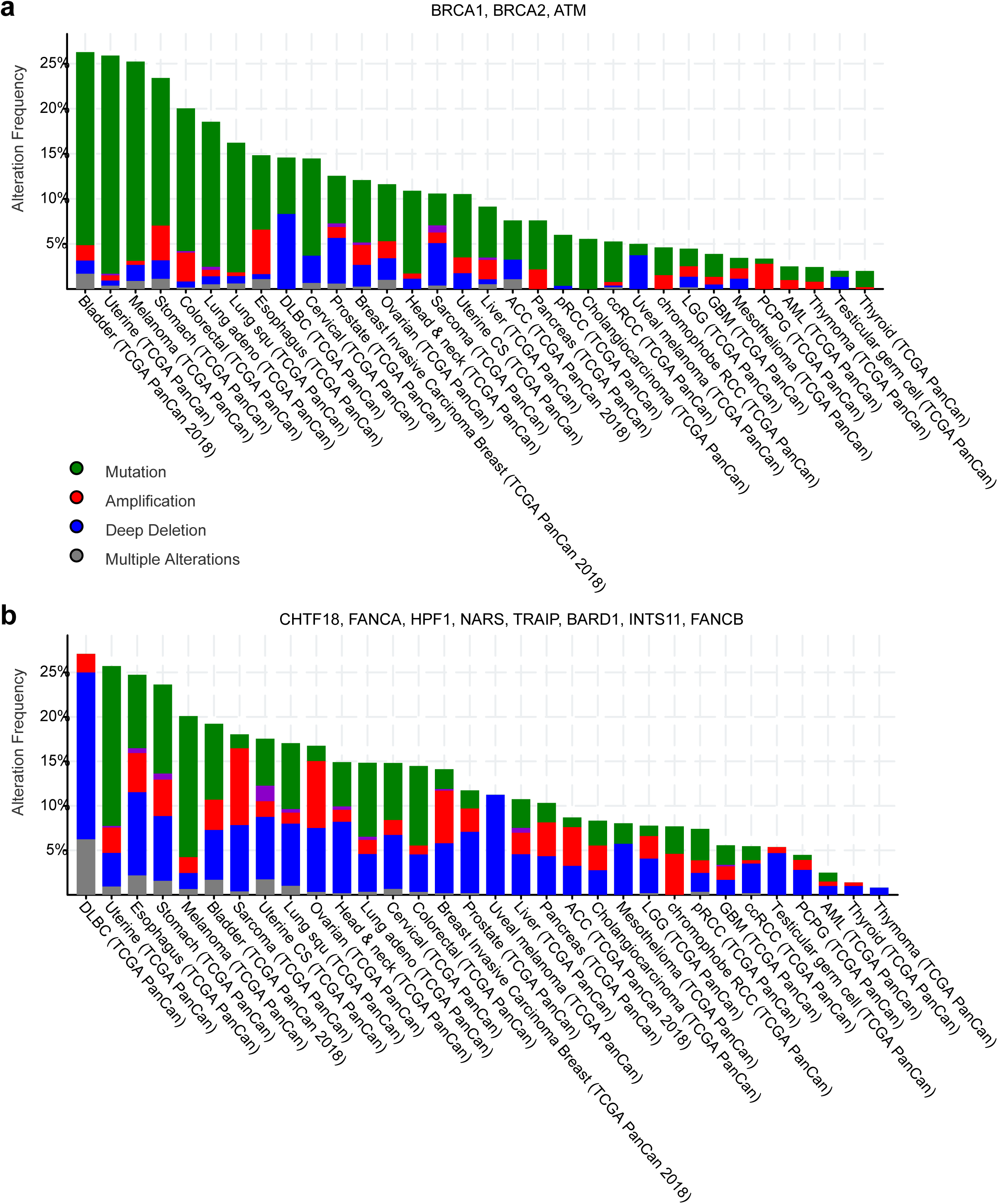
Alteration Frequency plots from cBioPortal for PARP-related genes. The TCGA PanCancer Atlas Studies were queried via the web interface for these genes, and the resulting plots are shown here. (a) Combined frequencies for BRCA1, BRCA2, and ATM. (b) Combined frequencies for CHTF18, FANCA, HPF1, NARS, TRAIP, BAD1, INTS11, and FANCB.

